# Genomic profiling of childhood tumor patient-derived xenograft models to enable rational clinical trial design

**DOI:** 10.1101/566455

**Authors:** Jo Lynne Rokita, Komal S. Rathi, Maria F. Cardenas, Kristen A. Upton, Joy Jayaseelan, Katherine L. Cross, Jacob Pfeil, Laura E. Ritenour, Apexa Modi, Alvin Farrel, Gregory P. Way, Nathan M. Kendsersky, Khushbu Patel, Gonzalo Lopez, Zalman Vaksman, Chelsea Mayoh, Jonas Nance, Kristyn McCoy, Michelle Haber, Kathryn Evans, Hannah McCalmont, Katerina Bendak, Julia W. Böhm, Glenn M. Marshall, Vanessa Tyrrell, Karthik Kalletla, Frank K. Braun, Lin Qi, Yunchen Du, Huiyuan Zhang, Holly B. Lindsay, Sibo Zhao, Jack Shu, Patricia Baxter, Christopher Morton, Dias Kurmashev, Siyuan Zheng, Yidong Chen, Jay Bowen, Anthony C. Bryan, Kristen M. Leraas, Sara E. Coppens, HarshaVardhan Doddapaneni, Zeineen Momin, Wendong Zhang, Gregory I. Sacks, Lori S. Hart, Kateryna Krytska, Yael P. Mosse, Gregory J. Gatto, Yolanda Sanchez, Casey S. Greene, Sharon J. Diskin, Olena Morozova Vaske, David Haussler, Julie M. Gastier-Foster, E. Anders Kolb, Richard Gorlick, Xiao-Nan Li, C. Patrick Reynolds, Raushan T. Kurmasheva, Peter J. Houghton, Malcolm A. Smith, Richard B. Lock, Pichai Raman, David A. Wheeler, John M. Maris

## Abstract

Accelerating cures for children with cancer remains an immediate challenge due to extensive oncogenic heterogeneity between and within histologies, distinct molecular mechanisms evolving between diagnosis and relapsed disease, and limited therapeutic options. To systematically prioritize and rationally test novel agents in preclinical murine models, researchers within the Pediatric Preclinical Testing Consortium are continuously developing patient-derived xenografts (PDXs) from high-risk childhood cancers, many refractory to current standard-of-care treatments. Here, we genomically characterize 261 PDX models from 29 unique pediatric cancer malignancies and demonstrate faithful recapitulation of histologies, subtypes, and refine our understanding of relapsed disease. Expression and mutational signatures are used to classify tumors for *TP53* and *NF1* inactivation, as well as impaired DNA repair. We anticipate that these data will serve as a resource for pediatric oncology drug development and guide rational clinical trial design for children with cancer.

**Highlights:**

- Multiplatform genomic analysis defines landscape of 261 pediatric cancer patient derived xenograft (PDX) models
- Pediatric patient derived xenografts faithfully recapitulate relapsed disease
- Inferred *TP53* pathway inactivation correlates with pediatric cancer copy number burden
- Somatic mutational signatures predict impaired DNA repair across multiple histologies

## Introduction

An estimated 15,780 children and adolescents (< 20 years) are diagnosed with cancer in the United States each year, and these diverse entities are the leading cause of disease-related deaths in children (American Childhood Cancer Organization 2014). Despite five-year survival rates for pediatric cancers now exceeding 80%, survivors frequently have lifelong side effects from cytotoxic therapy, and survival outcomes for children with certain types of tumors remain dismal. The relative rarity of pediatric cancers, molecular and mechanistic heterogeneity of subtypes within and across histologies, genetic and molecular distinction from adult malignancies, tumor evolution in the face of cytotoxic standard therapies, and lack of targeted therapeutic agents all pose major challenges to improving outcomes for children with cancer. Indeed, there are very few drugs with specific labeled indications for pediatric malignancies, and the majority of standard therapies are largely empiric.

Preclinical testing of new therapeutic anti-cancer agents is essential in the field of pediatric oncology due to the relative rarity of the condition and the need to prioritize agents for early phase clinical trials. Over the past 15 years, the Pediatric Preclinical Testing Consortium (PPTC), previously known as the Pediatric Preclinical Testing Program (Houghton et al. 2007; Houghton et al. 2002) has developed over 370 patient-derived xenograft (PDX) models from high-risk childhood cancers. In collaboration with pharmaceutical and academic partners, the PPTC systematically screens novel therapeutic agents for anti-tumor efficacy in order to help prioritize those that will move to the clinic. While some of these models have been credentialed with mRNA expression arrays (Whiteford et al. 2007) and/or single nucleotide polymorphism (SNP) arrays (El-Hoss et al. 2016), here we present a comprehensive genomic characterization of 261 models from 29 unique pediatric cancer malignancies.

## Results

### Analysis pipeline for somatic mutations, gene expression, RNA fusions, and copy number profiling in pediatric PDX tumors

We performed whole exome sequencing (WES) on 240 childhood cancer PDX models, whole transcriptome sequencing (RNA-Seq) on 244 models, single nucleotide polymorphism (SNP) microarrays on 252 models (Table S1, Figure S1 panels A-B), and performed short tandem repeat (STR) profiling on all 261 models (Figure 1B, Table S2). Of the 261 models profiled, 82 had available references that are also included in Table S2.

**Figure 1.**
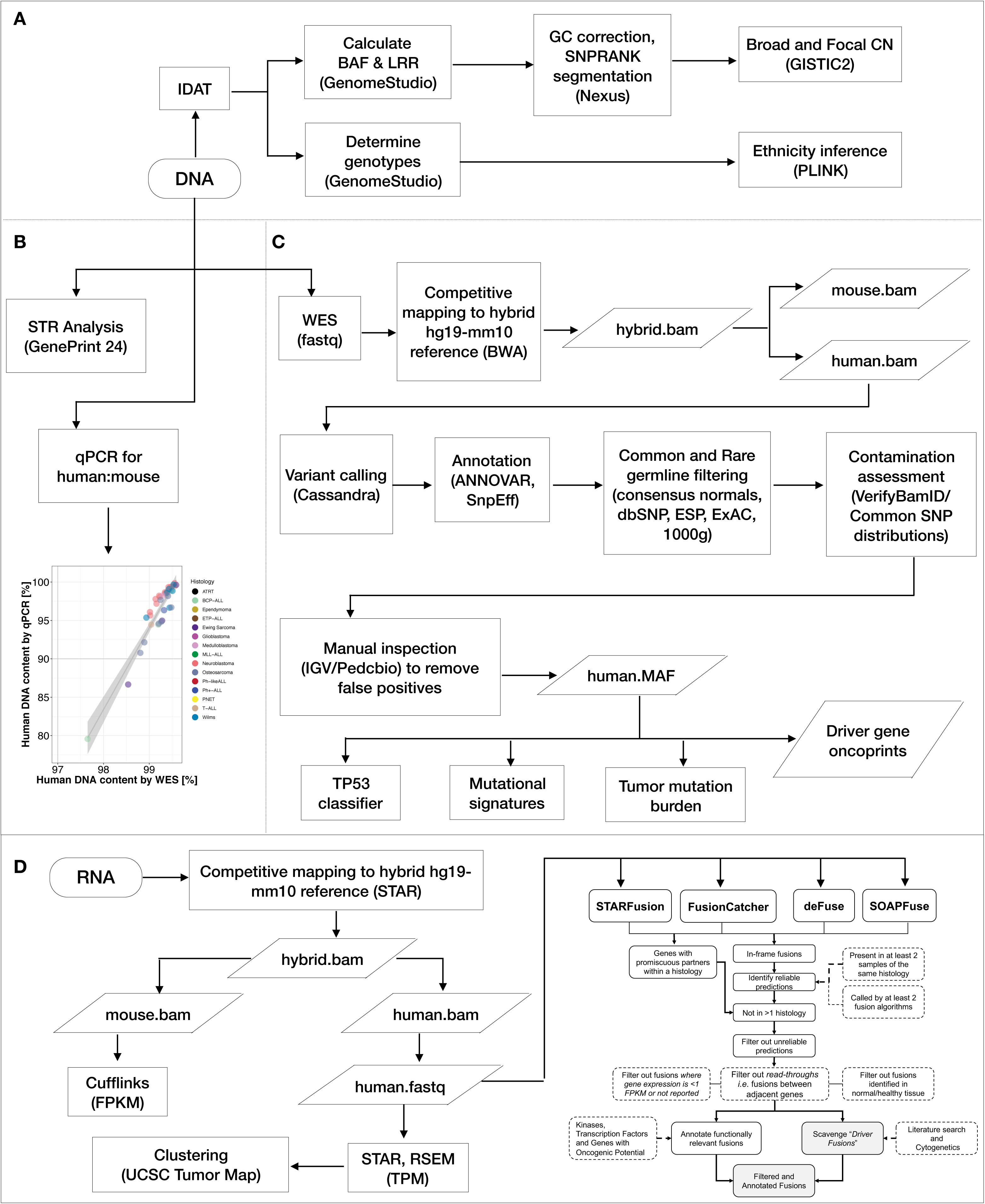
Analysis pipeline for somatic mutations, gene expression, RNA fusions, and copy number profiling in pediatric PDX tumors. Figure 1 displays an overview of analysis methods utilized. Genomic DNA from PDX tumors was used for SNP array copy number analysis (A, N = 252), short-tandem repeat identity testing (B, N = 261), quantitative PCR to assess human:mouse DNA content (B, N = 35), and whole exome sequencing (C, N = 240). Total RNA from PDX tumors was used for whole transcriptome sequencing (D, N = 244). See Table S1 for Ns per assay per histology and Table S2 for STR profiles.

Figure 1 depicts the analysis workflow (see STAR methods for details). Of the 240 models on which WES was performed, 69 models were previously sequenced through efforts of the PPTP (dbGAP ID phs000469.v17.p7), and we harmonized these data. For WES (Figure 1C) and RNA-Seq (Figure 1D), we performed competitive mapping to a hybrid human-mouse reference (hg19-mm10) and used human-specific BAM files as the bases for downstream analyses. We validated this biochemically with quantitative PCR (qPCR) by calculating the ratio of human:mouse DNA in a subset of 35 PDX tumors. We found a significant correlation between the percent of human reads following WES hybrid mapping and the percent of human DNA in the tumor extract (Figure 1B, Pearson correlation R = 0.943, F = 272.5, df = 34, p-value < 2.2e-16). A MAF file of common germline variation was created if a variant was present in more than five normal samples from TCGA patients (N = 809). The remaining variants, comprised of both somatic and rare germline alterations, were collated into the “somatic” MAF file. Artifactual sequencing variants were removed as described in STAR methods (Table S3). Mutation variation was summarized (Figure S1, panels C-I and Table S4). Common germline SNP distributions (allele frequency > 0.005 in any one of the three databases: Exome Aggregation Consortium, 1000 genomes, or the NHBLI Exome Sequencing Project) were plotted for each model and visually inspected for a negatively skewed distribution to assess DNA cross-contamination in WES data. To identify potential mis-identification, RNA variant calling was performed and variant allele frequencies correlated between WES and RNA. Models whose variants did not correlate were deemed mis-identified and removed (STAR Methods). Within this cohort, five pairs of models were derived from tissue at a single phase of therapy (Table S1). Thus, as additional QC, we correlated somatic mutation allele frequencies between each pair and found high concordance of mutation frequencies (Figure S1, panel J), confirming biological reproducibility of creating PDX models within a center. The OS-36/OS-36-SJ pair were derived from lung metastases at two different centers and interestingly, the OS-36 model contains a higher mutational load (117 total mutations) than the OS-36-SJ model (37 total mutations) which could reflect tumor heterogeneity in metastatic disease (Figure S1, panel K).

SNP arrays were processed for segmentation, focal copy number, and ethnicity inference (STAR methods, Figure 1A, Figure S2). As reported ethnicities were only available for a small proportion of the models, we used SNP array genotypes to infer approximate ethnicities using HapMap genotype frequencies. We assigned models to African, East Asian, European, and South Asian/Hispanic ethnicities (Figure S2, Table S1). Overall, 71% of models are of predicted European descent, 11.5% South Asian/Hispanic, 9.1% African, 5.5% mixed or unknown ethnicity, and 2.4% East Asian.

Following rigorous assessment for contamination, mis-identification, and sample mislabeling, 26 full models were removed and 3 RNA samples were removed. All remaining models used in these analyses were shown to be free of detectable levels of DNA contamination (STAR Methods).

### PDX models recapitulate the mutation and copy number landscape of childhood cancers

We defined oncogenic driver genes for broad histology classifications (brain, sarcoma/carcinoma, leukemia, renal, neuroblastoma, and osteosarcoma) in Table S5, based on previously reported pediatric cancer genomic studies (Ma et al. 2018; Gröbner et al. 2018; Pugh et al. 2013; Eleveld et al. 2015; Behjati et al. 2017; Shern et al. 2014; Liu et al. 2017; Zhang et al. 2012). RNA fusions, somatic mutations, and focal copy number events for the top 20 altered genes are presented in Figure 2, full oncoprint matrices in Table S5, and described below for samples on which at least WES was performed. Focal homozygous deletions correspond to loss of expression (FPKM < 1) in models for which RNA was available. Genome-wide copy-number profiles for histologies with 10 or more models are plotted in Figure S4. *ATRX* deletions were determined from WES bam files (STAR Methods). Pathway enrichment derived from RNA-Seq for histologies with N >=4 are plotted in Figure 6C.

**Figure 2.**
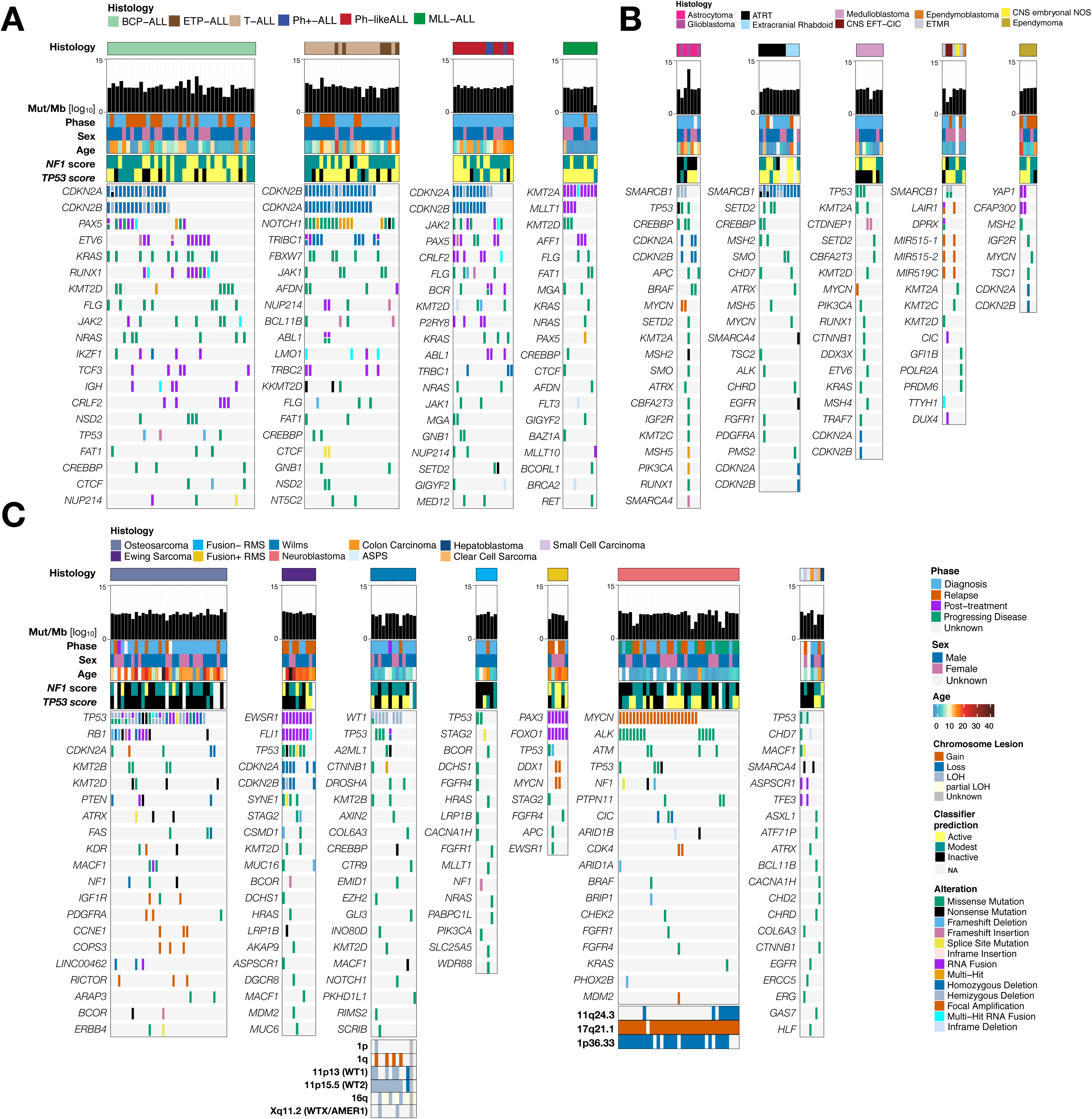
Patient-derived xenograft models recapitulate the mutational landscape of childhood cancers. Oncoprints of somatic alterations (homozygous deletions, amplifications, SNVs, and fusions) in known driver genes for PDX models for which exome sequencing was performed (N = 240, top 20 genes per histology shown, or fewer if total < 20). Oncoprints are grouped by acute lymphoblastic leukemia (A: From left to right are B-cell precursor ALLs (N = 37), early T-cell precursor (ETP) and T-cell ALLs (N = 25), Philadelphia chromosome-like (Ph-like) and Philadelphia chromosome positive (Ph+) ALLs (N = 18), and mixed lineage leukemias (MLL, N = 10)), central nervous system (B: From left to right are: astrocytic tumors, comprised of astrocytoma (N = 4) and glioblastoma (GBM, N = 3), atypical teratoid rhabdoid tumor (ATRT, N = 8) and extracranial rhabdoid tumors (N = 4), medulloblastomas (N = 8), non-medulloblastoma embryonal tumors comprised of embryonal tumor with multilayered rosettes, C19MC-altered (ETMR, N = 3), CNS Ewing sarcoma family tumor with CIC alteration (CNS EFT-CIC, N = 2), ependymoblastoma (N = 1), and CNS embryonal tumor not-otherwise specified (CNS embryonal tumor NOS, N = 1), and ependymomas (N = 5)), and extracranial solid tumors (C: From left to right are osteosarcomas (N = 34), Ewing sarcomas (N = 10), Wilms tumors (N = 13), fusion negative rhabdomyosarcomas (N = 6), fusion positive rhabdomyosarcomas (N = 6), neuroblastomas (N = 35), and rare tumors (N = 7)). Clinical annotations for all models include: histology, patient phase of therapy from which PDX was derived, sex, and age. Additional histology-specific annotations include: *TP53* classifier score for osteosarcoma models, 1p, 1q, 11p13, 11p15.5, 16q, and Xq11.2 status for Wilms tumor models, and 1p36.33, 11q24.3, and 17q21.2 status for neuroblastoma models. Hemizygous deletions in *TP53* are annotated for osteosarcoma models, in *CDKN2A* for leukemia models, and in *WT1* for Wilms tumor models. See Table S2 for driver gene lists. For BCP-ALL, T/ETP-ALL, Ph+/-like-ALL, and osteosarcomas, only mutations observed in more than one model are depicted.

#### Acute Lymphoblastic Leukemias

Figure 2A depicts oncoprints for 90 acute lymphoblastic leukemia models.

##### BCP ALLs

A total of 41-43% of BCP-ALL PDX models contain canonical focal deletions of the tumor suppressors on chromosome 9p, *CDKN2A* or *CDKN2B* (Figures 2A and S4B), the majority of which are homozygous. ALL-07 and ALL-19 harbor both a *CDKN2A* homozygous deletion and a *CDKN2B* hemizygous deletion. ALL-84 has hemizygous loss of both *CDKN2A/B* and contains a nonsense mutation in, and minimal expression of *CDKN2A* (FPKM = 3.14). The BCP-ALL models were enriched for alterations in the RAS pathway (*KRAS* mutated in 27%, *NRAS* mutated in 16%), the JAK-STAT pathway (*JAK2/3* altered in 22%), and 22% have altered *KMT2D*. These pathways, along with *PI3K/AKT, TNFα,* and *TP53* signaling, were all significantly enriched in the expression data (Figure 6C). We detected fusion transcripts in 29 BCP-ALL models (78%); 24% with *ETV6* fusions (66% partner with *RUNX1*), 14% with *PAX5* fusions, 14% with *TCF3* fusions, and 8% with *IKZF1* fusions (Table 1).

**Table 1.**
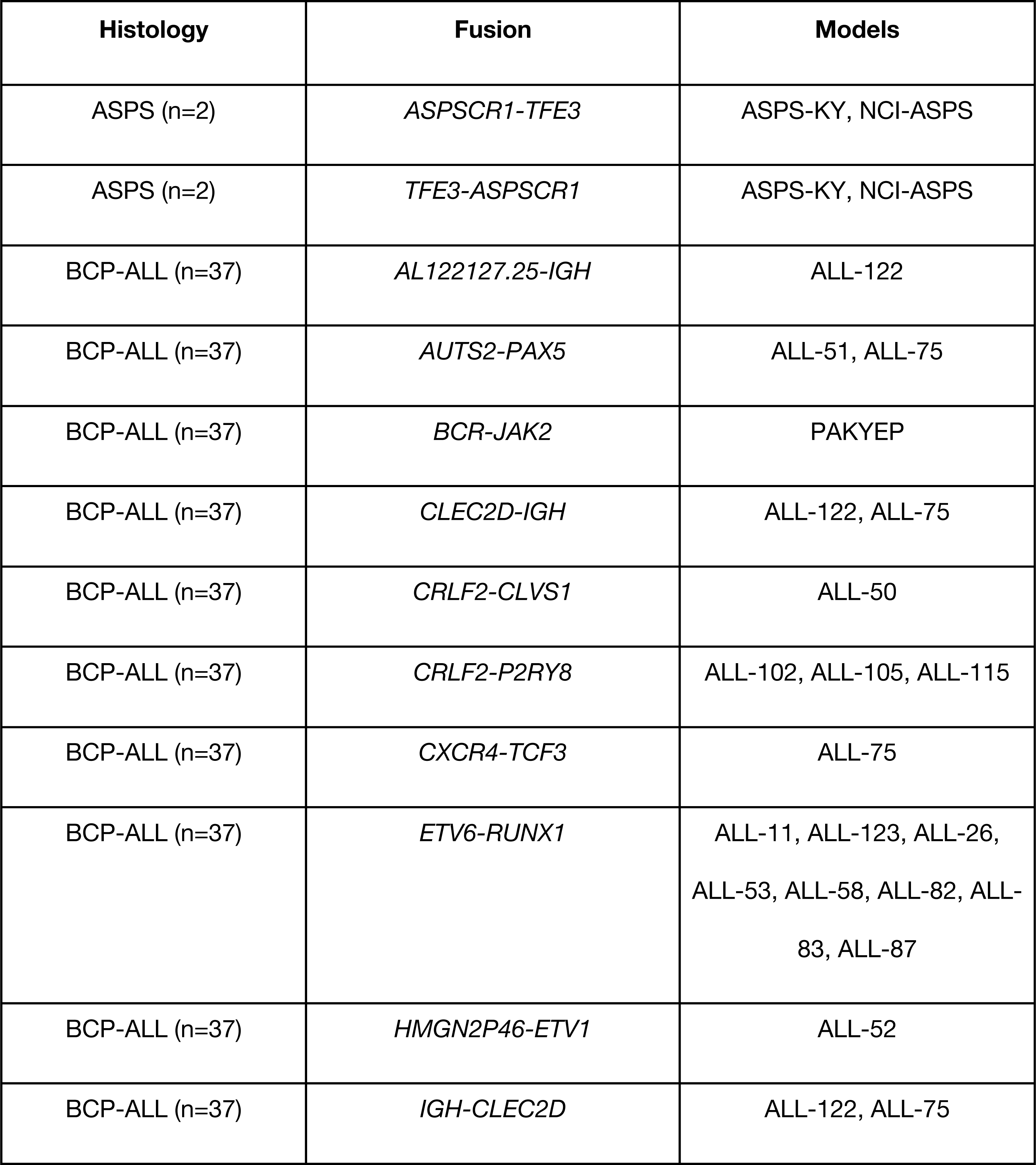
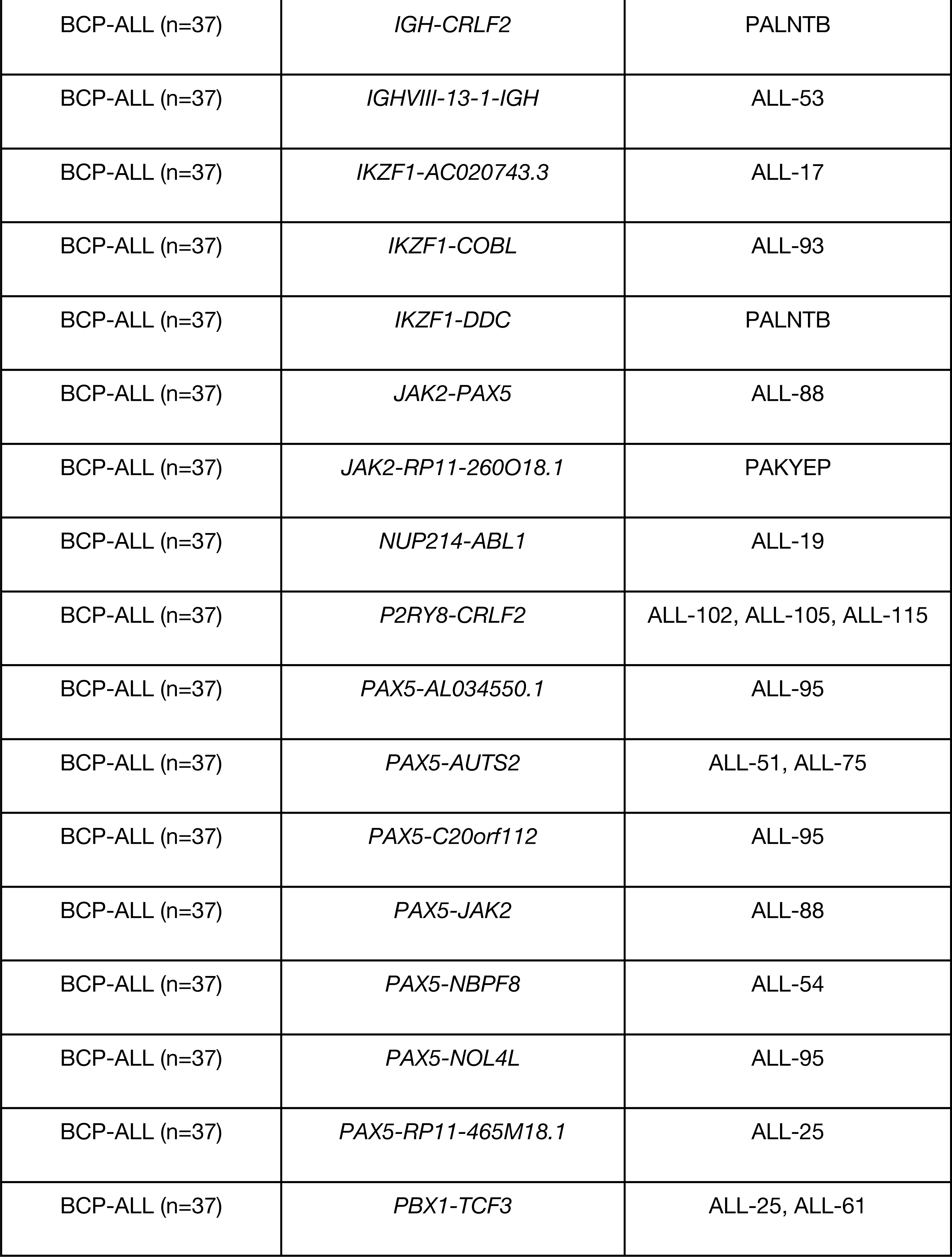
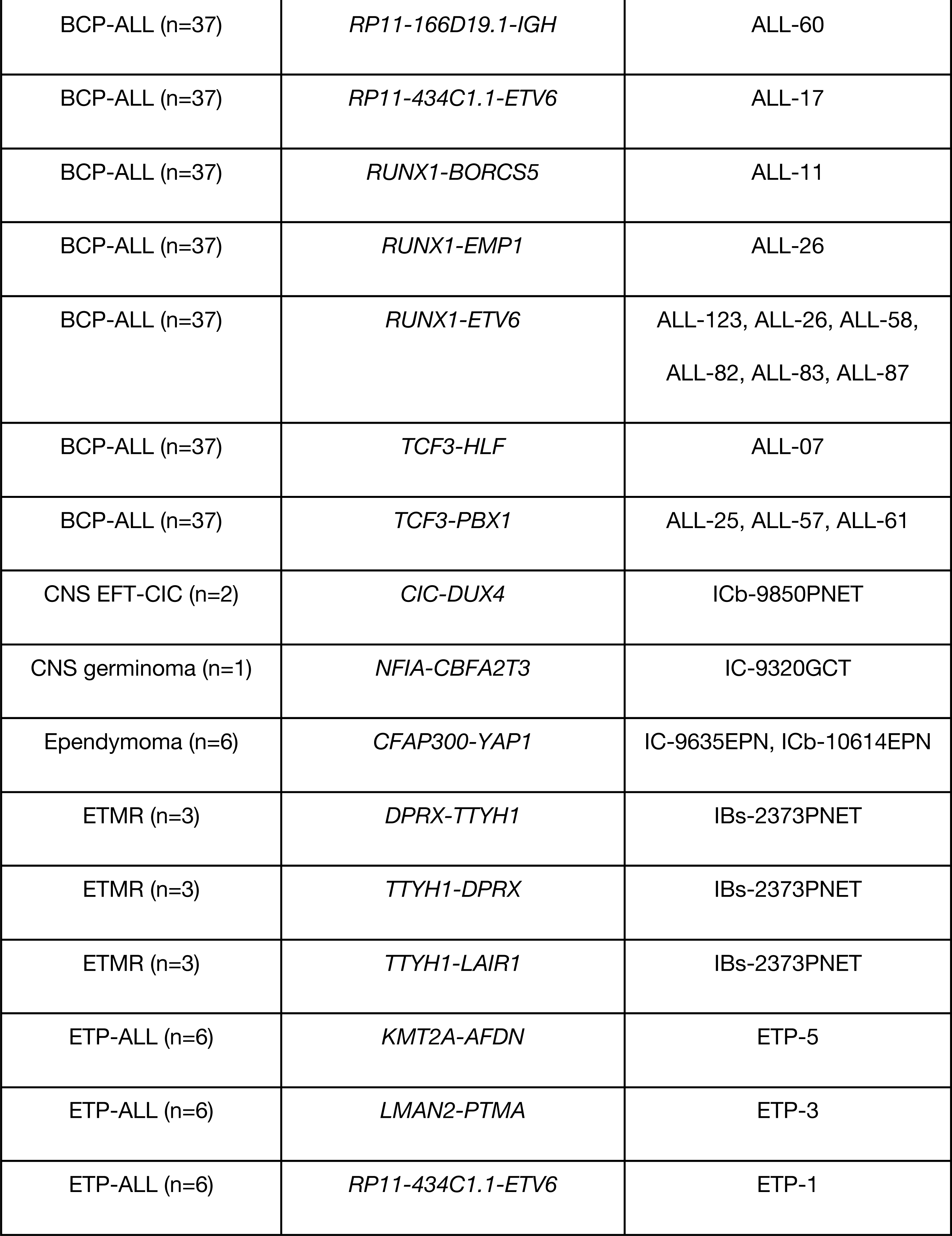
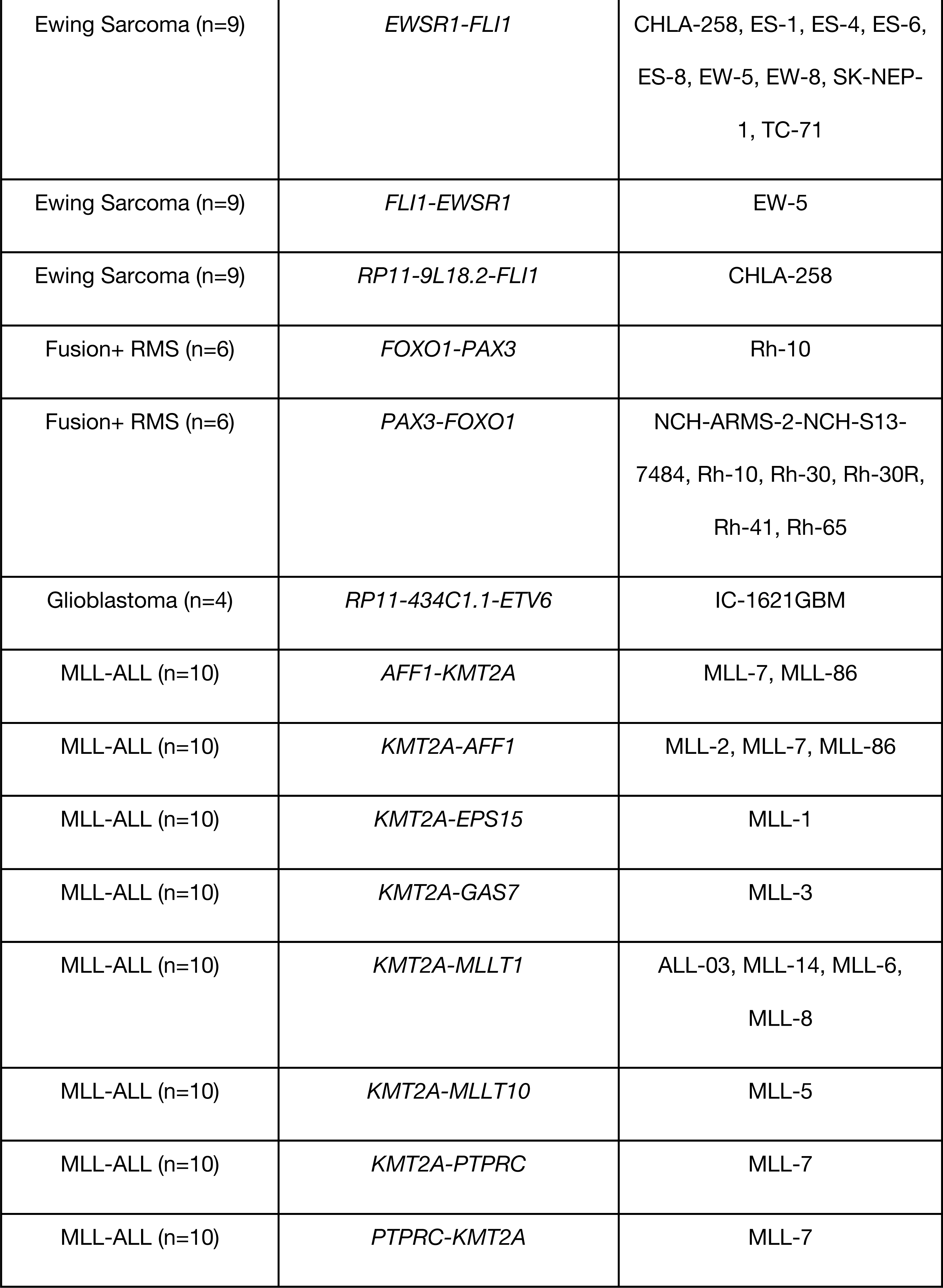
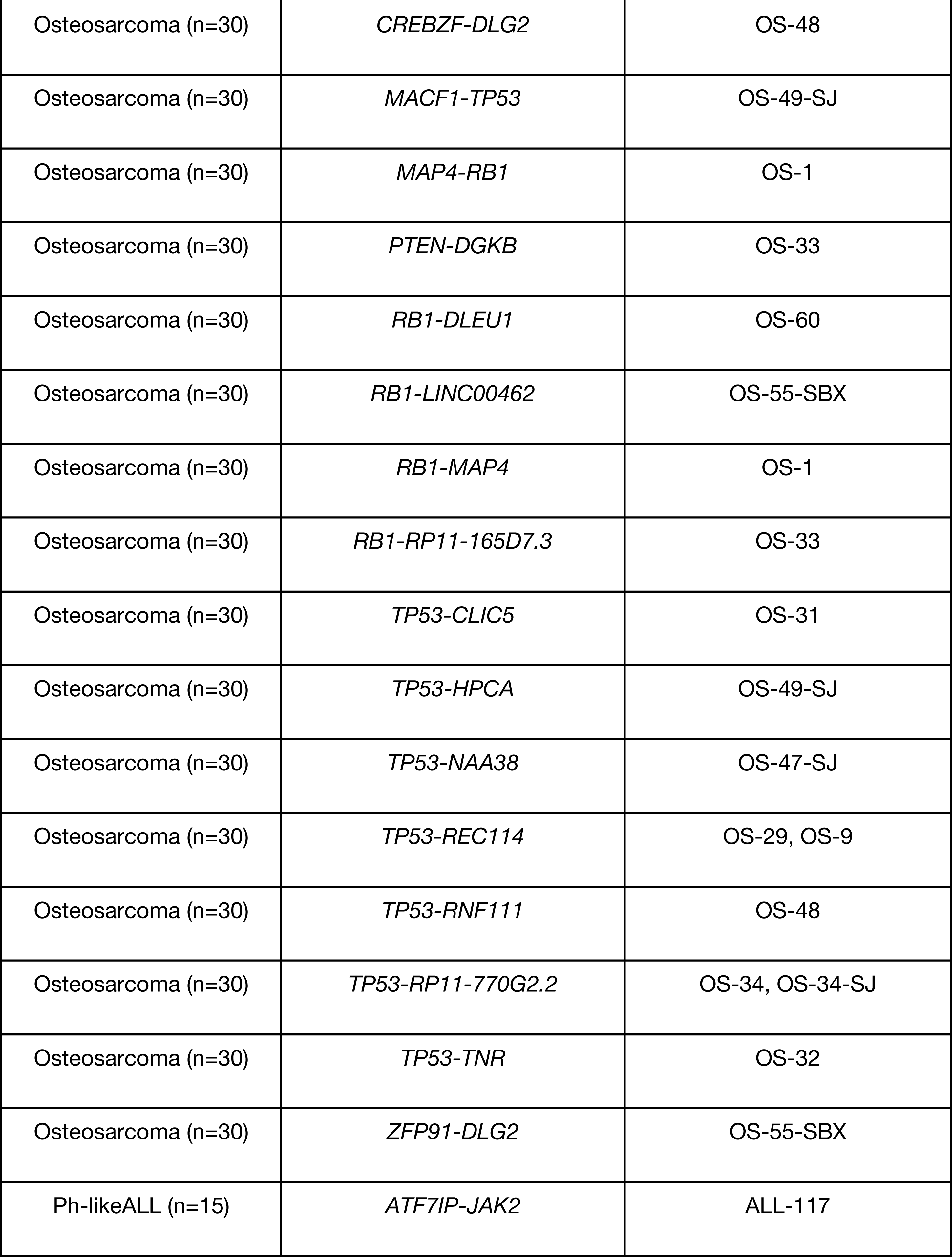
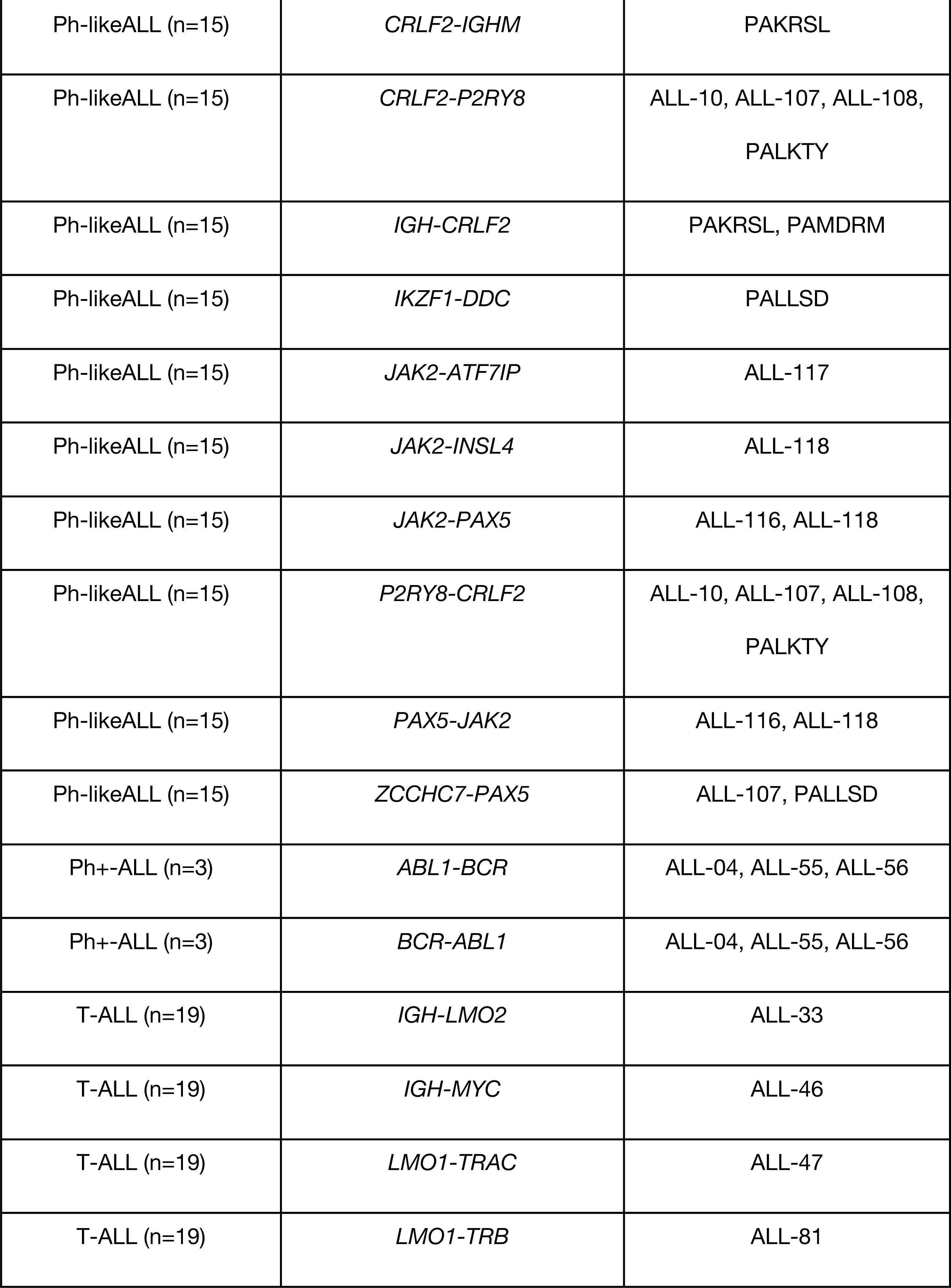
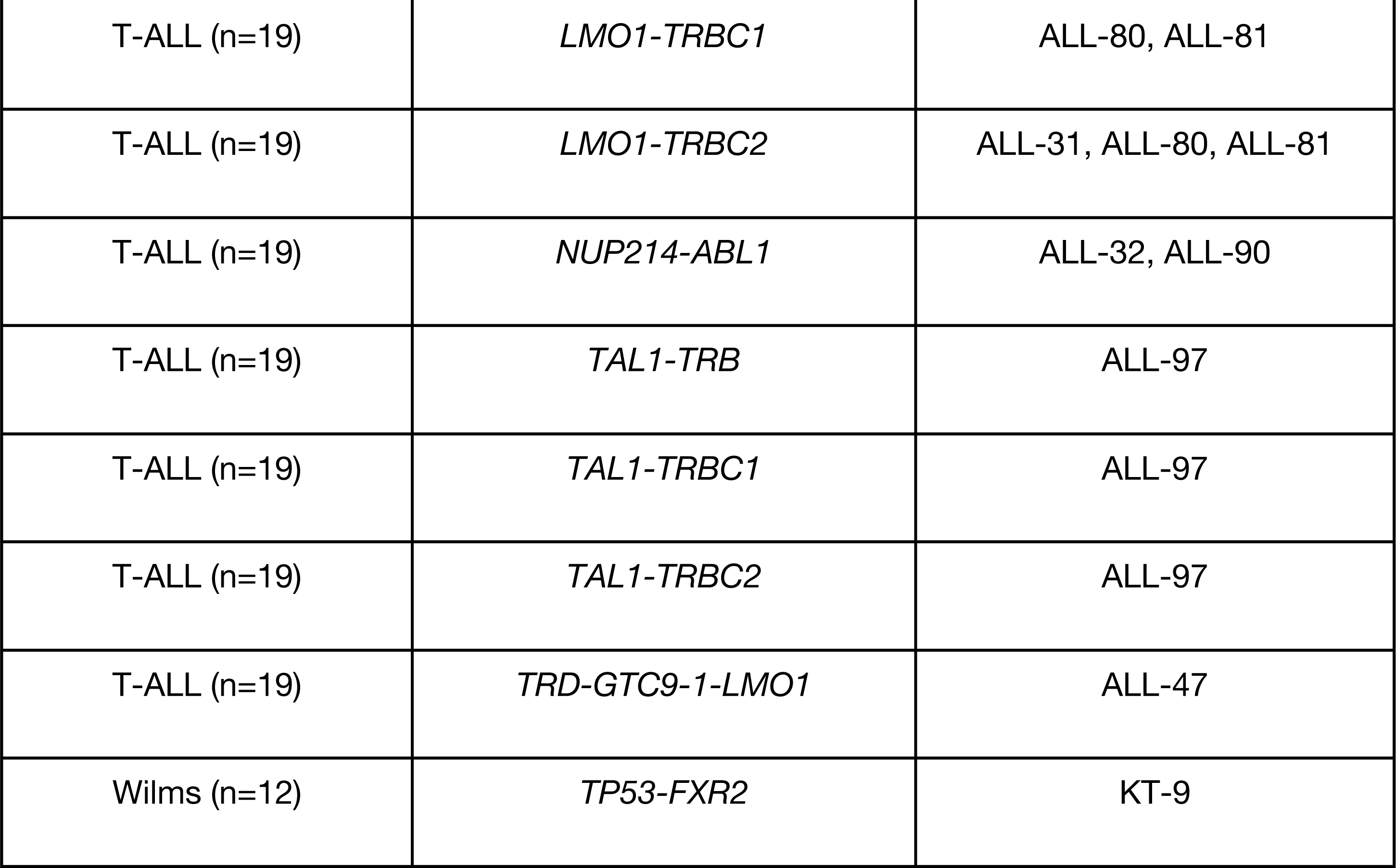
Fusions in known driver genes that were detected in this PDX dataset. Following rigorous filtering (Figure 1D, STAR Methods), 92 unique fusions in 101 unique models were found in known histology-specific driver genes.

##### ETP and T-ALLs

ETP-ALL and T-ALL models are predominantly characterized by *CDKN2A/B* focal deletions (72-76%, Figure S4B) and/or a *NOTCH1* mutation (68%). ALL-43 has a focal homozygous deletion of *CDKN2B* and partial homozygous deletion of *CDKN2A* but retains mRNA expression of *CDKN2A* (FPKM = 12). Genes within the JAK-STAT pathway are also frequently altered: lesions in *JAK1 or JAK2* are observed in 24% of models and 4% of models contain lesions in *STAT5B.* We detected oncogenic fusion transcripts in nearly half (48%) of these models involving the following genes: *TRBC2* (16%), *TRBC1* (12%), *ABL1* (8%), *IGH* (8%), *LMAN2* (4%), *LMO1* (4%), *LMO2* (4%), and *ETV6* (4%). *ATRX* is deleted in ETP-1 and expression is ablated (FPKM < 5, Figure S3, panel A).

##### Ph-like and Ph+ ALLs

We confirmed a *BCR-ABL1* fusion in all three Ph+-ALL models (ALL-04, ALL-55, and ALL-56). Five of the Ph-like ALL models (33%) contain a canonical *CRLF2-P2RY8* fusion. Additional frequently-rearranged genes include *JAK2* (13%) *and PAX5* (27%). In both Ph+ and Ph-like ALL models, focal deletion of CDKN2A/B (56-67%, Figure S4B) and IKZF (44%) alterations are predominant. Frequently-altered pathways include Ras and JAK-STAT (Figure 2A, 6C).

##### MLL-ALLs

All MLL-ALL models contain a canonical *KMT2A* fusion and have relatively silent genomes (Figure 2A) with minimal copy number alterations (Figure S4B). The majority of these models were derived from children < 1 year of age.

#### Central nervous system and extracranial rhabdoid tumors

In Figure 2B, we present the mutational spectrum of PDX models derived from CNS and extracranial rhabdoid tumors.

##### Glial-derived models

Of the three GBM tumors in this cohort, IC-1621GBM harbored a pathogenic histone gene missense mutation (*HIST1H3A* D78N) as well as a *TP53* missense mutation. This PDX was generated from a patient with DNA mismatch repair deficiency syndrome and showed 124 somatic mutations per MB. We confirmed multiple mutations in mismatch repair genes *PMS1, MSH2, MSH5,* and *POLE* (non-exonuclease domain mutation). The likely oncogenic drivers are the nonsense mutations in *PMS1* (Q316*) and *MSH2* (G721*), each of which disrupt the DNA mismatch repair protein domain and the MutS domain, respectively (Figure S3, panel B). IC-6634GBM harbors a hemizygous *SMARCB1* mutation, missense mutation in *TP53,* amplification of *MYCN*, and focal deletion of *CDKN2A/B.* NCH-PXA-1 and IC-3635PXA were derived from pleomorphic xanthoastrocytomas, one of which harbors a canonical *BRAF* V600E point mutation. NCH-MN-1 was derived from a patient diagnosed with an anaplastic rhabdoid meningioma with the clinical suspicion of an ATRT, however this model had no evidence of an inactivating *SMARCB1* alteration. Rather, it harbors a *BRAF* V600E mutation and focal *CDKN2A/B* deletion, classifying this model as a high-grade glioma, herein denoted as an astrocytoma. IC-2664PNET was derived from a patient diagnosed with a primitive neuroectodermal tumor (PNET), but was further molecularly classified as a high-grade astrocytoma. IC-2664PNET has focal amplification of *MYCN* and a hemizygous *SMARCB1* deletion, but retains mRNA expression of *SMARCB1* (Figure S2, panel C).

##### ATRT and extracranial rhabdoid tumors

All eight ATRT models and the four extracranial rhabdoid tumors harbor inactivating alterations (focal deletion, frameshift deletion, or nonsense mutation) in *SMARCB1*, the hallmark tumor suppressor gene deleted in rhabdoid tumors.

##### Medulloblastomas

Of the 21 medulloblastoma models, 10 could be classified into one of the four consensus molecular subgroups based on histopathology and genomic profiling (Northcott et al. 2017). Six models were classified as the SHH subtype: ICb-1078MB, ICb-1197MB, ICb-1338MB, ICb-2123MB, ICb-5610MB, ICb-984MB, one as WNT subtype: NCH-MB-1, two as Group 3 subtype: ICb-1494MB and ICb-1595MB, one as Group 4 subtype: ICb-1487MB, and 10 were unclassified (pathology unavailable and genetics inconclusive, Table S1). NCH-MB-1 was reported to be derived from a large cell anaplastic tumor, but this model harbors missense mutations in *TP53, CTNNB1,* and *DDX3X*, and has monosomy of chromosome 6, thus genetically classifying this model into the WNT subtype.

##### Non-medulloblastoma embryonal tumors

All tumors previously labeled clinically as PNETs were reclassified. ICb-S1129MB was classified as an embryonal tumor with multilayered rosettes (ETMR) by pathology and indeed has high expression of the ETMR biomarker, *LIN28A* (Figure S3, panel D). Copy number profiling was not performed on this tumor. ICb-1343ENB and IBs-2373PNET were genetically classified as ETMR due to amplification of C19MC and high expression of *LIN28A*. Additionally, IBS-2373PNET contains a *DPRX-TTYH1* reciprocal fusion and a *TTYH1-LAIR1* fusion. ICb-9850PNET and IC-22909PNET-rIII, a diagnosis/relapse pair, were genetically classified as CNS EFT-CIC, as the diagnostic tumor contains a *CIC-DUX4* fusion. These models also harbor hemizygous *SMARCB1* deletions and could be differentially classified as ATRT, though both retained some mRNA expression of *SMARCB1* (Figure S3, panel C). The previously-annotated PNET model BT-27 was genetically classified as CNS embryonal NOS. The ependymoblastoma model IC-1499EPB did not harbor known driver mutations.

##### Ependymomas

Of the six total ependymoma models, one contained a *C11orf95-RELA* fusion IC-1425EPN (no WES; not shown in oncoprint) and two contained a *CFAP300-YAP1* fusion, IC-9635EPN and ICb-10614EPN (no WES; not shown in oncoprint), classifying these three as supratentorial.

##### DIPGs

The two DIPG models were profiled with RNA-Seq and SNP array, thus are not shown in the oncoprint. Both IBs-P1215DIPG and IBs-W0128DIPG show expression of *H3F3A* and *H3F3B* (FPKM > 50), genes encoding the histone H3.3 variant, and lack expression of *HIST1H3B* or *HIST1H3C,* genes encoding the histone H3.1 variant (Figure S3, panel E). However, we did not detect H3.1 or H3.3 histone mutations in these models. RNA variant calling revealed IBs-W0128DIPG contained predicted damaging (Polyphen) missense mutations in *NRAS* (p.G13R, 0.41), *CIC* (p.C102Y, 0.44), *SNCAIP* (p.R806H, 0.44), *KMT2C* (p.C988F, 0.45), *and PTCH1* (p.P1315L, 0.45). We did not detect any CNS driver gene damaging mutations in IBs-P1215DIPG.

#### Extracranial solid tumors

Figure 2C depicts the landscape of PDX models derived from extracranial solid tumors.

##### Osteosarcomas

The hallmark of osteosarcomas is *TP53* inactivation, and using a classifier trained on RNA expression data from The Cancer Genome Atlas (TCGA), we found all osteosarcoma models with available RNA-Seq data (N = 32) were predicted to have non-functional *TP53* (described below). As expected, *TP53* was the most commonly-altered gene (82%) in osteosarcoma PDX models (Figure 2C). Additional genes commonly altered include: *RB1* (35%), *CDKN2A* (15%), *KMT2B, KMT2D,* and *PTEN* (each 12%), and *ATRX* (9%). Of note, *ATRX* harbors a partial deletion in OS-41 (exons 2-26, FPKM = 11) and the full gene is deleted with ablation of expression in OS-45-TSV-pr1, and OS-60 (Figure S3, panel A). Recurrent focal amplifications were observed in *IGF1R, PDGFRA, CCNE1, COPS3, CDK4, RICTOR,* and *MYC* (6-9%). Osteosarcoma genomes demonstrate global copy-number changes, consistent with high prevalence of complex genomic rearrangements found in this tumor type (Figure S4).

##### Ewing sarcomas

The canonical *EWSR1-FLI1* fusion was found in all Ewing sarcoma models profiled with RNA-Seq (NCH-EWS-1 was not profiled) and CHLA-258 contained an additional *FLI1* fusion partner: *RP11-9L18.2* (Table 1, Figure 2C). *TP53* mutations are present in seven (70%) cases, with six showing allele frequencies at or near 1.0 due to copy-neutral loss of heterozygosity (cnLOH, ES-6, EW-8, SK-NEP-1) or loss of heterozygosity (LOH) due to chromosomal arm deletion (EW-5, ES-8, and TC-71), and the mutation in ES-1 appears hemizygous (R248Q, AF = 0.44) (Figure S3, panel F). Homozygous *CDKN2A/B* loss (60%) was mutually exclusive of *STAG2* mutations (20%), as expected (Tirode et al. 2014). We observe canonical (Tirode et al. 2014) broad gain of whole chromosomes 8 and 12, as well as focal 1q gain and 16q loss, in Ewing sarcomas (Figure S4A).

##### Wilms tumors

The mutational and copy number landscapes of Wilms tumor (N = 13) PDX models are depicted in Figures 2C and S4A. The *WT1* gene located at 11p13 was mutated in one PDX model (NCH–WT–6–S13–1506), but we observed hemizygous deletions of *WT1* in 61% of Wilms models, many of which were due to LOH of the entire 11p13 region. The 11p15.5 region, which contains imprint control regions (ICR) 1 and 2, often undergoes loss of imprinting (LOI) either due to maternal DNA methylation or maternal LOH/paternal uniparental disomy (pUPD) in Wilms tumor. The 11p15.5 region harbored LOH in 69% (9/13) of Wilms tumors, consistent with previous reports (Scott et al. 2012). Two models (15%) harbored hemizygous deletions of *AMER1* (formerly known as *WTX* and/or *FAM123B*), annotated as LOH of Xq11.2 (manual inspection; X and Y chromosomes were removed from copy number analyses). KT-9 is the only model annotated as coming from a patient with bilateral disease, and although it does not harbor a *WT1* mutation, it does have two hits in *TP53:* a *TP53-FXR2* fusion and a partial homozygous deletion. The *TP53* RNA fusion breakpoint was predicted to be in the 5’ UTR, which is concordant with SNP array data showing DNA deletion breakpoints within *FXR2* and *TP53* (Figure S3, panel G). Two Wilms models (15%, KT-6 and NCH-WT-6-S13-1506) harbored *CTNNB1* mutations, and consistent with previous reports (Scott et al. 2012), these were mutually exclusive of *WTX* alterations. Gain of the 1q arm, 1p LOH, and 16q LOH, adverse prognostic biomarkers for Wilms tumors (Segers et al. 2013; Spreafico et al. 2013; Pan et al. 2017), were observed in 31% (4/13), 8% (1/13), and 23% (3/13) of models, respectively. Additionally, KT-13, KT-10, and NCH-WT-4 had partial gains and/or mixed hyperploidy at 1q and KT-6, NCH-WT-4, KT-10, NCH-WT-5, and KT-8 had hotspots of LOH on 16q. Model KT-5 was not profiled with a SNP array.

##### Rhabdomyosarcomas

The oncoprints for fusion positive (Fusion+ RMS, N = 6) and fusion negative (Fusion- RMS, N = 6) rhabdomyosarcomas are depicted in Figure 2C. All Fusion+ RMS models harbored a hallmark *PAX3-FOXO1* fusion (Figure 2C, Table 1). Median patient age of Fusion+ RMS patients (16 years) was higher than that of Fusion-RMS patients (five years) (Table S4). As expected, we also observe focal amplification of *MYCN* (Rh-65 and NCH–ARMS–2–NCH–S13–7484) and *CDK4* (NCH–ARMS–2–NCH–S13–7484 and Rh-30). Amplification of *CDK4* was not present the Rh-30R (relapse tumor paired with Rh-30, SNPs and STRs confirm identity). Fusion-RMS models contained nonsynonymous mutations in previously reported recurrently-mutated genes at expected frequencies. For example, combined Ras pathway mutations (*NRAS, HRAS, KRAS, and NF1)* are typically observed in one-third of Fusion-RMS cases and here, Ras mutations were observed in 3/6 models (Rh-12 with *NF1 T2335fs,* NCH–ERMS–1–NCH–RMS–1 with *NRAS Q61K* mutation, and Rh-36 with *HRAS* Q61K). Other genes previously documented as recurrently-mutated in fewer than 10% of Fusion-RMS cases and found in this PDX dataset were: *FGFR4, PIK3CA, and BCOR.* All models except for IRS-68 over-express the common rhabdomyosarcoma biomarker, *MYOD1* (Figure S3, panel H). IRS-56, Rh-12, Rh-36, and Rh-70 had 11p15.5 LOH.

##### Neuroblastomas

Amplification of the *MYCN* oncogene was the most frequent alteration observed across all models (66%), and as expected, is largely mutually exclusive of 11q deletion (23%). Seventy seven percent of models had 1p deletion and 97% had 17q gain (collapsed profiles are shown in Figure S4A). Focal amplifications were observed in *CDK4* (6%), *FGFR4* (3%), and *MDM2* (3%) and homozygous deletions in *CIC* (9%). Consistent with previous reports, we find *ALK* to be the most frequently mutated gene (37% of all models contain hotspot mutations) and neuroblastomas that progress (36%) or relapse (50%) contain a higher frequency of *ALK* mutations than PDX models derived from diagnostic material (25%). Missense, nonsense, indel, and/or splice, mutations were found in: *TP53* (11%), *ATM* (11%), *PTPN11* (9%), *NF1* (9), *ARID1B* (6%), *BRAF* (3%), *BRIP1* (3%), *CHEK2* (3%), *CIC* (3%), *FGFR1* (3%), *FGFR4* (3%), *KRAS* (3%), and *PHOX2B* (3%). The nonsense and frameshift deletions in *NF1* correspond with ablated expression in COG-N-590x and NB-1771, respectively, but NB-1643 retains expression. COG-N-625x, derived from a male child, harbors an *ATRX* deletion (exons 2-10) (Figure S3, panel A).

##### Rare histologies

Seven PDX models were derived from rare tumor types and are depicted in Figure 2C: hepatoblastoma (NCH-HEP1), alveolar soft part sarcoma (ASPS-KY and NCI-ASPS), colon carcinoma (NCH-CA-3), small cell carcinoma, large cell variant (NCH-CA-1 and NCH-CA-2, diagnosis and relapse pair), and a clear cell sarcoma (NCH-CCS-1). Three models (43%) contained alterations in *TP53*; of note, an in-frame hemizygous deletion of *TP53* evolved at relapse in NCH-CA-2. The canonical *ASPSCR1-TFE3* fusion was detected in both ASPS models. NCH-CA-1 and NCH-CA-2 harbored deleterious *SMARCA4* mutations and NCH-CA-3 harbored a deleterious *NF1* nonsense mutation, each with concurrent loss of mRNA expression and as such, may be potential drivers of oncogenesis in these tumors. NCH-HEP1 contained a likely oncogenic WNT pathway mutation (*CTNNB1* p.D32G).

#### Breakpoint Density

We calculated total number of breakpoints per sample and breakpoint density within chromosomes, the latter as a surrogate measure of putative chromothripsis events (STAR methods). Consistent with pediatric cancer genomics literature, we observe very few breakpoints per sample in hematologic malignancies compared to those in solid tumors (mediaN = 3 breakpoints per sample in CNS embryonal NOS to mediaN = 154.5 breakpoints per sample in osteosarcoma; Figure S4C and Table S4). We found 25% (64/252) of models profiled have high breakpoint density (HBD) across one or more chromosomes (Figure S4D, Table S4), consistent with a recent pan-cancer chromothripsis report (Cortes-Ciriano, 2018). Specifically, 97% of osteosarcomas had HBD; 30% (10/33) of these contained HBD on four or fewer chromosomes indicative of chromothripsis, while the remaining 70% (23/33) contained HBD on five or more chromosomes, supporting a more complex and chaotic genomes prevalent in this tumor type (Lorenz et al. 2016). In neuroblastoma samples, 17% of models contained HBD on chromosomes 2, 5, 16, 17, 19. Chromothripsis events on chromosomes 2, 5, and 17 in neuroblastoma tumors have been previously reported associated with *MYCN* amplification, *TERT* rearrangements, and 17q gain, respectively (Molenaar et al. 2012; Boeva et al. 2013). Recurrent loci with HBD in medulloblastoma were chromosomes 2, 8, 14, and 17, consistent with recent reports (Rausch et al. 2012). In summary, PDX models faithfully recapitulate important prognostic copy number alterations of pediatric tumors.

### TMB and clonal evolution

Using somatic missense and nonsense mutations, we calculated tumor mutation burden (TMB) for each PDX model (see STAR methods). Median TMB across all models was 2.66 somatic mutations per megabase (Mut/Mb; Figure S4, panel E and Table S4). The TMBs across this cohort of PDX models are likely higher than those in previous reports for two main reasons. First, 37% of the PDX models were derived from a patient tumor at a phase of therapy other than diagnosis (Figure S1, A) and it is now known that tumors acquire significantly more somatic mutations post-therapy and following a relapse (Eleveld et al. 2015; Schramm et al. 2015; Padovan-Merhar et al. 2016; Ma et al. 2015; Schleiermacher et al. 2014). In fact, we observe an overall significantly higher TMB in PDX models derived from relapse tissue (3.08 Mut/Mb) compared to those derived at diagnosis (2.57 Mut/Mb, Wilcoxon p-value = 0.002, Figure 3A). Histologies containing models derived from both diagnosis and relapse are plotted in Figure S4, panel F. Second, without paired normal samples, rare germline and private variants could not be reliably removed from the “somatic” MAF. Thus, TMB reported here is likely inflated, but the trends across histologies and phase of therapy should accurately reflect TMBs determined with a paired germline sample.

**Figure 3.**
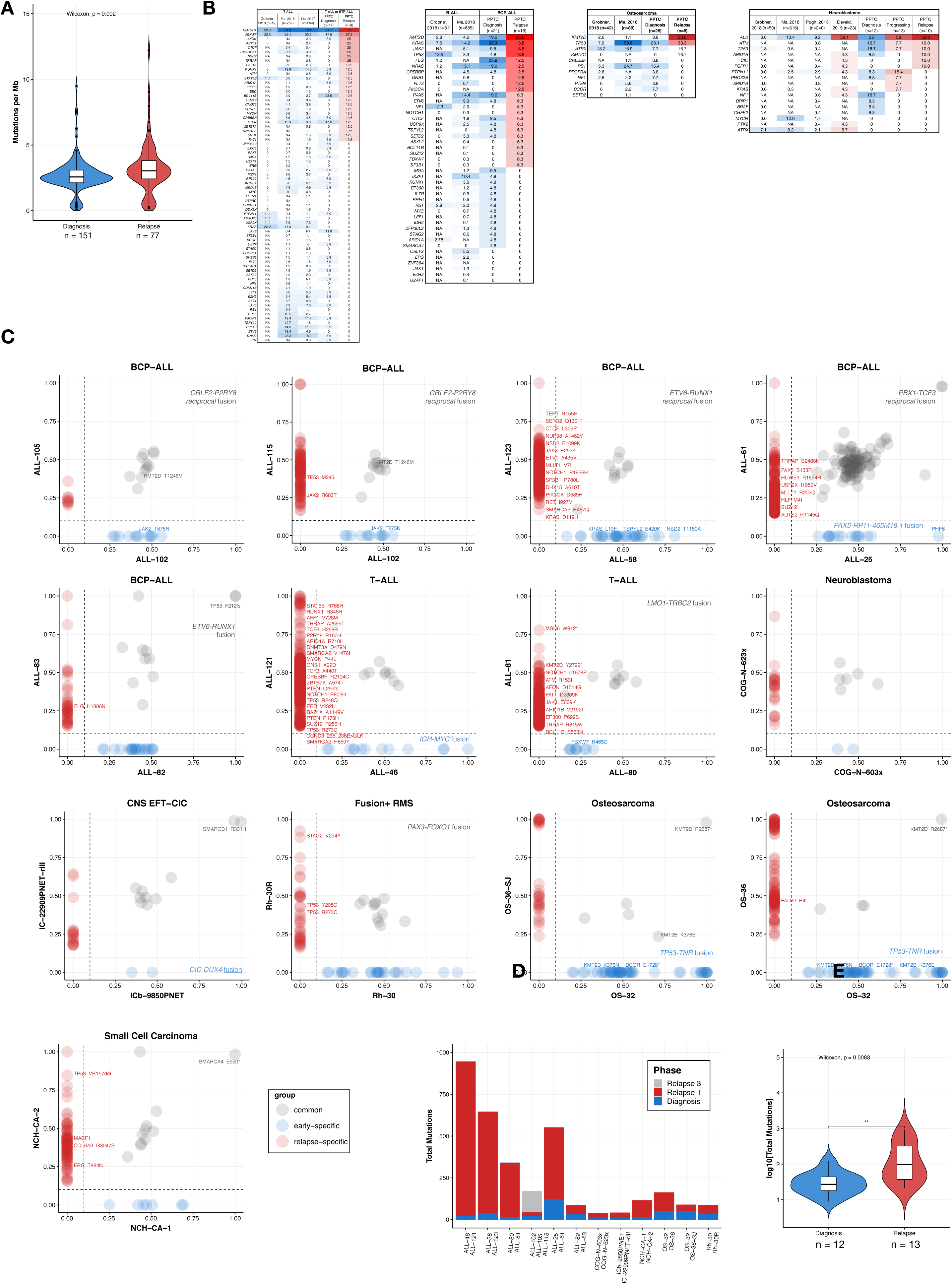
TMB and clonal evolution. Models derived from relapsed tumors contain a significantly higher number of mutations compared to models derived from diagnostic tumors (A, Wilcoxon p = 0.002, N_diagnosis_ = 151, N_relapse_ = 77). Panel B depicts frequencies of SNVs within genes of interest for primary and relapse T-ALL (N_diagnosis_ = 17, N_relapse_ = 8), BCP-ALL (N_diagnosis_ = 21, N_relapse_ = 16), osteosarcoma (N_diagnosis_ = 25, N_relapse_ = 6), and neuroblastoma (N_diagnosis_ = 12, N_progressing_ = 13, N_relapse_ = 10) models compared to literature frequencies reported for primary tumors (blue = diagnosis, red = relapse, intensity scaled to highest value within each phase and histology; copy number variation and fusions not available for each source and not included). Frequency of genes altered within the primary PPTC cohort are concordant with primary tissue results reported in the literature and frequencies of major driver genes increase in relapse tumors (T-ALL: *NOTCH1*; BCP-ALL: *KMT2D, JAK2, TP53, ETV6*; Osteosarcoma: *TP53, ATRX, IGF1R;* Neuroblastoma: *ALK, PTPN11, ATM, TP53*). Panel C depicts scatterplots of mutant allele frequencies for matched diagnosis/relapse models (blue = early-specific mutations, grey = mutations common at diagnosis and relapse, red = late-specific mutations). Only mutations for genes of interest per histology and fusions are annotated. There increase in mutational burden upon relapse in each pair (D) was significant (E, Wilcoxon p = 0.0083, N_diagnosis_ = 12, N_relapse_ = 13).

The majority of the PDX models were established at diagnosis (63%), but 6% were derived from surgical resection specimens after neoadjuvant therapy, 27% from a relapsed specimen (14% of those were neuroblastomas from a large volume blood draw obtained immediately after death from disease progression), and 4% did not have phase of therapy annotated. In addition, 12 pediatric cancer patients had either two or three models created across the spectrum of their therapy (Table S1). The tables in Figure 3B report non-silent mutation frequencies for genes of interest (Table S5) as reported previously (Ma et al. 2018; Gröbner et al. 2018; Pugh et al. 2013; Liu et al. 2017). We report PDX gene mutation frequencies for histologies with paired diagnosis/relapse cohorts with group N ≧ 6: BCP-ALL (N_diagnosis_ = 21/N_relapse_ = 16), T-ALL or ETP-ALL (N_diagnosis_ = 17/N_relapse_ = 8), osteosarcoma (N_diagnosis_ = 25/N_relapse_ = 6), and neuroblastoma (N_diagnosis_ = 12/N_progressing_= 13/N_relapse_ = 10). Across all four histologies, there is an increased frequency of driver gene mutations in relapsed disease as indicated numerically and by the intensity of the red heatmap compared to those at diagnosis (blue heatmap). Next, we plotted somatic mutation allele frequencies for the 13 PDX diagnosis-relapse pairs (Figure 3C, 3D) and highlight diagnosis-specific (blue), relapse-specific (red), common (grey) mutations, demonstrating the increased frequency of relapse-specific mutations. Finally, the median number of mutations in the paired relapse samples was significantly higher than the number of mutations detected in the diagnostic samples (Figure 3E; median of 98.0 vs. 27.5 mutations; Wilcoxon p < 0.01).

### Expression and mutational signatures classify pediatric PDX models for TP53 inactivation, NF1 inactivation, and defective DNA repair

A recent study used TCGA data to classify tumors for *TP53* inactivation status and found that alterations in multiple genes phenocopy *TP53* inactivation, indicating that *TP53* mutation status alone is not necessary to infer inactivation of the pathway (Knijnenburg et al. 2018). We used a machine learning algorithm to infer *TP53* inactivation, as well as *NF1* inactivation and Ras pathway activation, from transcriptomes of PDX tumors using classifiers previously trained on TCGA expression data (STAR methods) (Knijnenburg et al. 2018; Way et al. 2018; Way et al. 2017). The *TP53* (AUROC = 0.89) and *NF1* (AUROC = 0.77) classifiers are both accurate compared to a shuffled gene expression baseline, but performance of the Ras classifier (AUROC = 0.58) was relatively poor (Figure 4A). Classifier scores > 0.5 predict inactivation of *TP53* or *NF1* (Table S6) and *TP53* scores are significantly higher (Wilcoxon p < 2.2e-16) in models with a *TP53* alteration (mean score = 0.747) compared to those without alterations (mean score = 0.459) (Figure 4B). Many models annotated as wild-type *TP53* have high *TP53* inactivation scores (Figure 4B). We found models with alterations in genes such as *MDM2* and *RB1* also have high *TP53* inactivation scores. These alterations may phenocopy *TP53* alterations (Figure 4C, genes chosen as primary or secondary interactors of *TP53* defined by the TP53 KEGG signaling pathway). In Figure 4D, we plot alterations for each gene by variant classification. Notably, all types of alterations within *TP53* were associated with high classifier scores, while scores for other genes varied by type of alteration.

**Figure 4.**
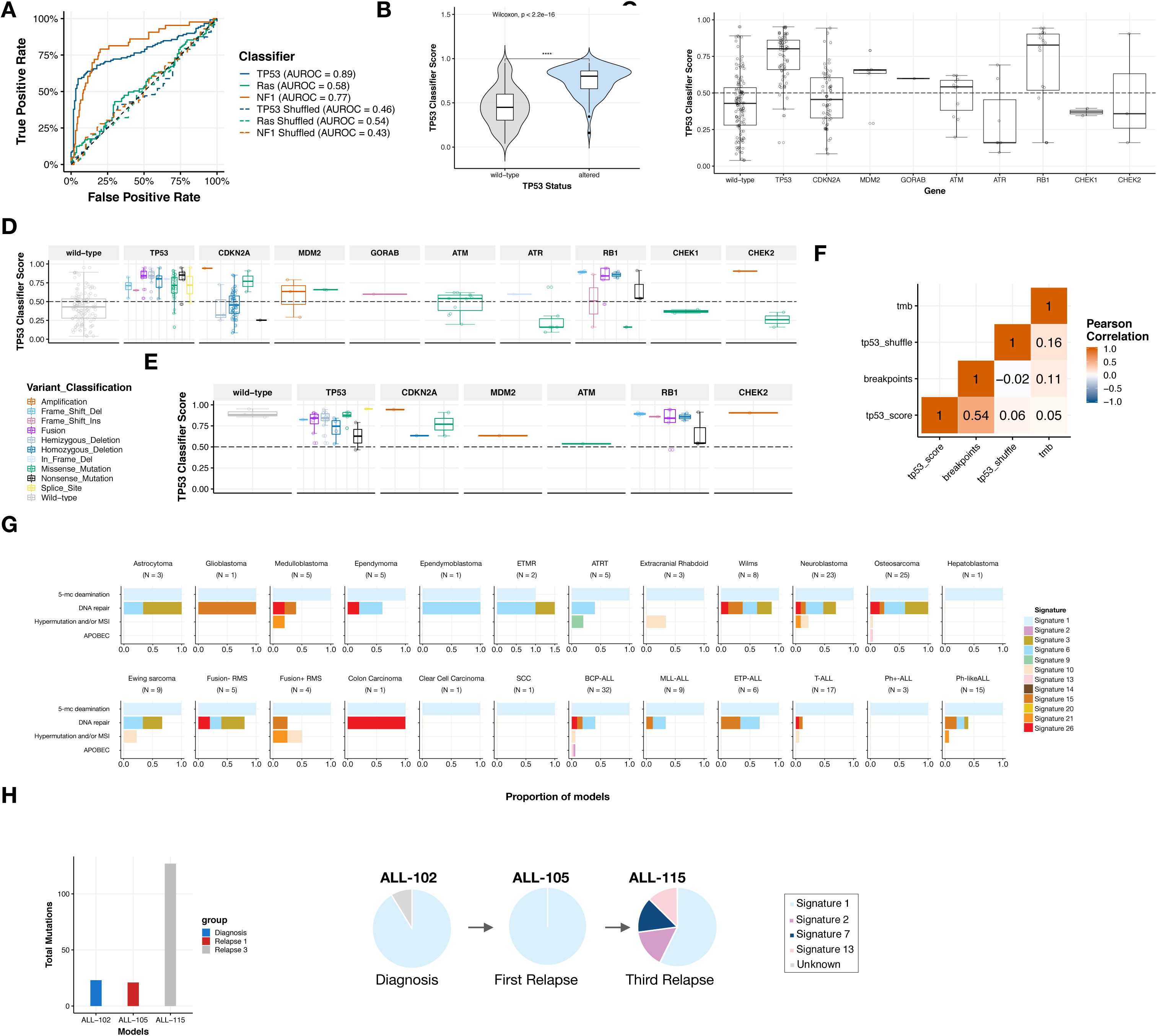
Expression and mutational signatures classify pediatric PDX models for TP53 inactivation, NF1 inactivation, and defective DNA repair. Only *TP53* and *NF1* classifiers performed well in our dataset (A, AUROC_TP53_ = 0.89, AUROC_NF1_ = 0.77, AUROC_Ras_ = 0.58). B, *TP53* scores are significantly higher (Wilcoxon p < 2.2e-16) in models with genetic aberrations in *TP53* (mean score = 0.747) compared to those without alterations (mean score = 0.459). C, Classifier scores are plotted by *TP53* pathway gene aberration and variant classification (D). In osteosarcoma models, most scores, regardless of variant type or gene, were predicted active (E). Overall copy number burden (number of breakpoints calculated from SNP array data, STAR methods) correlates significantly with *TP53* classifier score (F, R = 0.54, p = 2.0e-19). G, Somatic mutational signatures (STAR methods) for models containing > 50 mutations are plotted as cumulative barplots for signatures defined by 5-methylcytosine deamination (Signature 1), impaired DNA repair (Signatures 3, 6, 15, 20, and 26), hypermutation and/or microsatellite instability (Signatures 9, 10, 14, and 21), or APOBEC (Signature 2 and 13). Panel H displays an increased mutational load (total mutations) upon the third relapse (N_diagnosis(ALL-102)_ = 23, N_relapse1(ALL-105)_ = 21, N_relapse3(ALL-115)_ = 127). Panel I shows signatures 2 (APOBEC), 7 (UV exposure), and 13 (APOBEC) arose between the first and third relapse.

As *TP53* inactivation is a hallmark of osteosarcoma, we focused on these models as a proof-of-concept. The classifier predicted that all models profiled with RNA-Seq except OS-55-SBX had *TP53* pathway inactivation. Many had a genetic alteration in a *TP53* pathway gene as supporting evidence (Figure 4E, Table S6). However, the mechanisms of *TP53* inactivation in OS-34-SJ, OS-43-TPMX, and OS-51-CHLX are still unknown, and may require whole genome sequencing to detect. To ensure osteosarcoma models were not driving the observed association with *TP53* scores, we removed osteosarcoma models and reanalyzed the data. We found a significantly higher *TP53* classifier score (Wilcoxon p = 4e-12) in models with alterations in *TP53* pathway genes (Figure S5, panels A-B). We then evaluated which types of variants were associated with high *TP53* classification score and observed that models containing fusions had the highest classifier scores compared to wild-type, followed by models with SNVs and CNVs, respectively (Figure S5, panel C, Kruskal-Wallis P = 9.8e-11). These are broken down by gene in Figure S5, panel D. Outside of osteosarcomas, only one model contained a fusion in the *TP53* pathway: Wilms model KT-9 contained a *TP53-FXR2* fusion. We found overall copy number burden (number of breakpoints calculated from SNP array data, STAR methods), but not tumor mutation burden nor shuffle score, correlates significantly with *TP53* classifier score (Figure 4F, R = 0.54, p = 2.0e-19), supporting recent published observations (Knijnenburg et al. 2018). Genetic alterations rendering *TP53* inactive may contribute to copy number instability in these models. Use of gene expression classifiers can guide preclinical studies, for example, therapeutically targeting the *TP53* pathway in tumors with high *TP53* inactivation scores rather than those with altered *TP53*.

Next, we used the COSMIC 30 mutational signature profile to create single sample mutational signature matrices for each PDX model (Table S6, STAR methods). We validated the co-occurrence of Signatures 2 and 13 (R = 0.60, p = 8.18e-25) and independence of other DNA repair and hypermutation signatures from each other (Figure S5, panel E). Signature 1 (spontaneous deamination of 5-methylcytosine) was significantly inversely correlated with presence of defective DNA repair Signatures 3 and 6 (R = −0.41, p = 3.94e-11 and R = −0.54, p = 1.08e-19, Figure S5, panel E). Figure 4G displays signatures for models containing > 50 mutations and signatures with a cosine similarity > 0.1. In all brain tumor models, Wilms, neuroblastoma, osteosarcoma, Ewing sarcoma, both rhabdomyosarcoma subtypes, colon carcinoma, and all ALLs except for Ph+-ALL, we found models that contained signatures of defective DNA repair by homologous recombination (Signature 3) and/or DNA mismatch repair (Signatures 6, 15, 20, 26). Hypermutation (Signatures 9, 10, 14) and microsatellite instability (Signature 21) signatures were observed in 10 of 24 PDX histologies. The APOBEC mutational signature (Signatures 2 and 13) was found in two BCP-ALL models (ALL-58 and ALL-115) and one osteosarcoma model (OS-43-TPMX); both are histologies which have previously been reported to harbor this signature (Alexandrov et al. 2013). We did not observe kataegis in these models, but this may be a limitation of using WES instead of WGS (Figure S3, panel I). Figures 3C (first two plots) and 4H-I display tumor evolution across the one trio of models in this dataset: ALL-102 (diagnosis), ALL-105 (relapse 1), and ALL-115 (relapse 3). From diagnosis to the first relapse, there was little change in mutation burden. At diagnosis, ALL-102 contained an oncogenic *JAK2* T875N mutation (AF = 0.41), which appears to have been eradicated following standard chemotherapy (ANCHOG ALL8 trial ID ACTRN12607000302459); this clone was not detected in ALL-105 (first relapse). Upon the third relapse (ALL-115), total mutation burden significantly increased, *TP53* M246I (AF = 0.37, likely oncogenic) arose, and a new oncogenic *JAK2* mutation (R683T, AF = 0.26) arose (the second relapse was testicular and thus, sample was not available for engrafting), suggesting this tumor was driven by the *JAK2* and *TP53* oncogenic lesions. There was little change in mutational signature between diagnosis to the first relapse, but following the third relapse, additional Signatures 2 (APOBEC), 7 (UV exposure), and 13 (APOBEC) were detected (Figure 4I). The *CRLF2-P2RY8* reciprocal fusion persisted as the tumor evolved.

### Expression profiles of patient-derived xenograft models cluster by tissue of origin and contain driver fusions

We used the UCSC TumorMap (Newton et al. 2017) to visualize clusters of expression profiles across PDX histologies (Figure 5A). We observed clear separation among unrelated histologies and overlapping clustering among related histologies. For example, T-ALL and ETP-ALL cluster together as expected, but distinct from other ALL histologies. The leukemias clustered by subtype and distinctly from solid tumors. Ewing sarcoma, neuroblastoma, Wilms, and medulloblastoma form distinct clusters. Osteosarcomas cluster with two ASPS models. Fusion+ and fusion-RMS cluster near each other but distinctly. Brain tumor histologies cluster near each other with the exception of ATRTs, some of which cluster with extracranial rhabdoid tumors near sarcoma samples. We identified histology-specific expression differences using a Bayesian hierarchical model (Gelman, 2006), grouped related histologies under the same prior distribution and ranked gene expression differences for each histology and performed Gene Set Enrichment Analysis (GSEA). This demonstrated tissue-specific enrichment within each histology using GSEA and the TissGDB (Kim, 2018) and TiGER (Liu, 2008) gene sets (Figure 5B). To investigate pathway enrichment within histologies, we ran GSEA using the MSigDB curated (C2) gene sets and plotted the normalized enrichment scores (NES) for the Hallmark pathway gene sets in Figure 5C.

**Figure 5.**
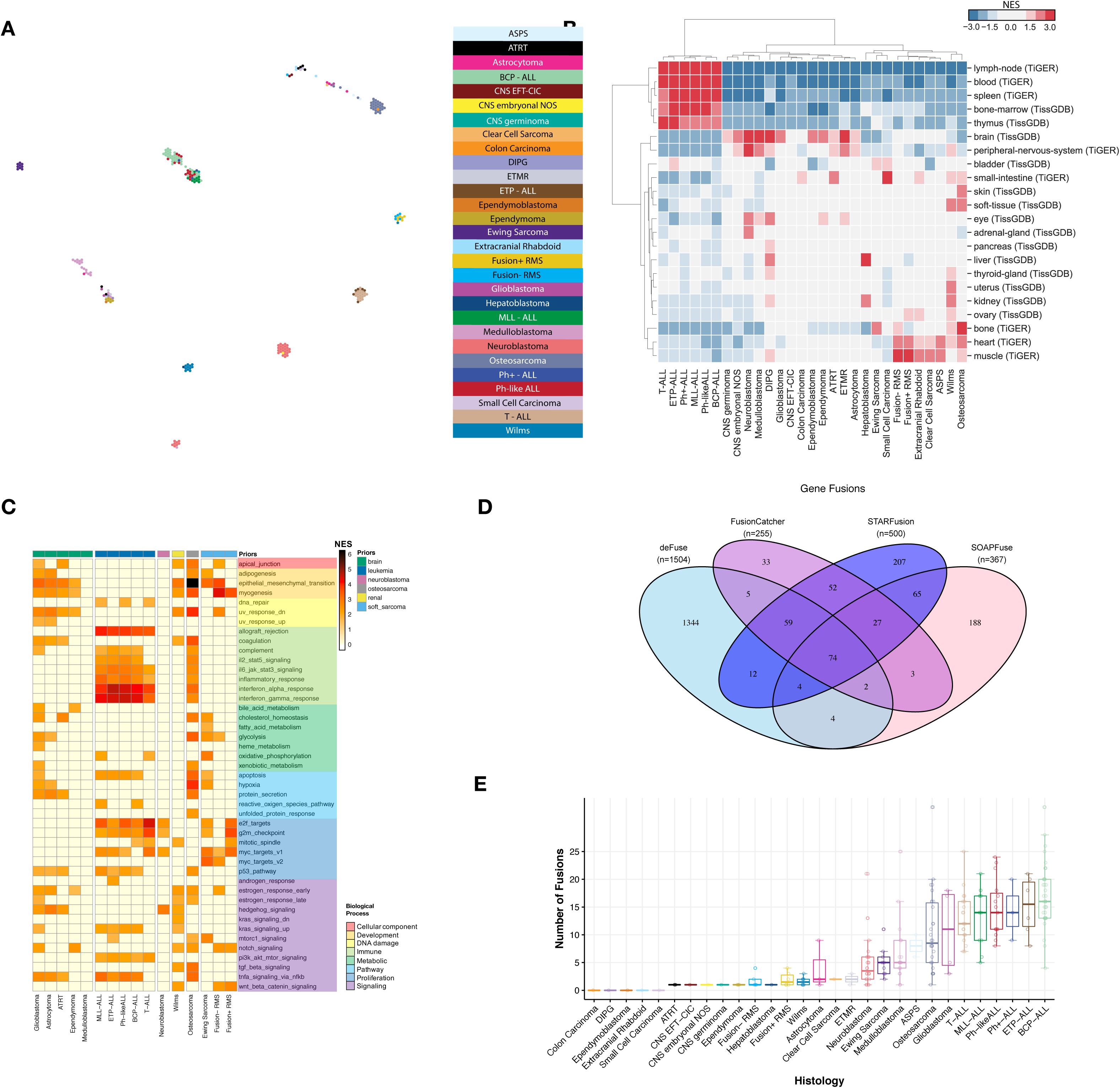
Expression profiles of patient-derived xenograft models cluster by tissue of origin and contain driver fusions. TumorMap rendition of PDX RNA-Seq expression matrices by histology (A) and hierarchical clustering depict tissue-specific enrichment within each histology (B, NES = normalized enrichment score). Gene set enrichment analysis for Hallmark pathways (C) demonstrate histology-specific biologic processes significantly altered (samples grouped by prior and process). Venn diagram of RNA fusion overlap among four algorithms (D) and high-confidence fusion totals (E) demonstrate a higher overall number of fusions in hematologic malignancies (boxplots are graphed as medians with box edges as first and third quartiles; detailed Ns in Table S4).

Next, we searched for fusions in the RNA-Seq data using four algorithms: defuse, FusionCatcher, STARFusion, and SOAPFuse. A total of 50,796 unique fusions were called and we used the pipeline described in Figure 1D and STAR methods to define 925 unique high-confidence fusions (Figure 5D and Table S7) and 92 unique known oncogenic driver fusions defined by cytogenetics and literature (Table 1 and Figure 5E). Fusions were annotated for frame and whether a gene partner is a known oncogene, kinase, or transcription factor to identify oncogenic potential and functional relevance. We found that PPTC PDX models largely maintain known oncogenic driver fusions specific to their histology: all alveolar rhabdomyosarcoma models harbored *PAX3-FOXO1* fusions, all Ewing sarcoma samples with RNA-Seq data showed *EWSR1-FLI1* fusions, all Ph+ ALL tumors contained *BCR-ABL1* fusions, and *KMT2A* (*MLL)* fusions were detected in all MLL-ALL models (Table 1). Osteosarcomas harbored *TP53* fusions and breakpoints reside within intron one of the *TP53* gene, a mechanism of TP53 inactivation which has been previously reported in osteosarcoma (Ribi et al. 2015). In five diagnosis/relapse pairs, we detected four fusions in the diagnostic PDX (*PAX5-RP11-465M18.1, IGH-MYC, CIC-DUX4,* and *TP53-TNR*) that were undetected in the paired relapse model (Figure 3C), suggesting these specific gene fusions may have been acquired after an alternative initiating event that was retained in each case.

## Discussion

In this study we used whole exome, whole transcriptome, SNP genotyping arrays, and STR profiling to genomically characterize 261 pediatric PDX models across 29 unique histologies. We developed novel analytical pipelines to remove mouse reads from DNA or RNA sequencing data and demonstrate high concordance between these pipelines and orthogonal measurement of human:mouse DNA. We validated faithful recapitulation of primary and relapsed disease within tumor of origin type through analysis of somatic mutations, copy number alterations, RNA expression, gene fusions, and oncogenic pathways.

The genomic and gene expression data presented herein have immediate applications to the prioritization of experimental agents for testing in pediatric preclinical models and in potentially moving them forward for clinical testing. For example, there are reports identifying specific genomic alterations as predicting sensitivity to ATR inhibitors, including ATM loss, ARID1A mutation, defective homologous recombination, and ATRX mutation associated with alternative lengthening of telomeres (ALT) (Lecona, 2018). Querying the PPTC data at PedcBioPortal can quickly identify models with these characteristics, and the models can then be used to test whether *in vivo* responsiveness to ATR inhibitors is predicted by one or more of the molecular characteristics. Similarly, PPTC RNA-seq data can be used to identify models that show elevated gene expression for the targets of immunotherapy agents such as antibody-drug conjugates and T cell engagers. As examples, in the PPTC dataset DLL3 shows overexpression in selected neuroblastoma and medulloblastoma models (Sano, 2018), ROR1 shows overexpression in a subset of B-ALL and Ewing sarcoma models, and IL3RA (CD123) shows overexpression in many B-ALL models. The PPTC RNA-seq dataset was also used to identify T-ALL as a target histology for an agent activated by the aldo-keto reductase AKR1C3 (Lock, 2018) and to identify alveolar soft part sarcoma (ASPS) xenografts as intrinsically overexpressing CD274 (PD-L1), making ASPS a target histology for the evaluation of checkpoint inhibition (OSullivan, 2018).

Further, we performed machine learning to classify tumors into *TP53* and *NF1* active or inactive and we suggest that these scores might be future biomarkers for drug response. These classifiers have been used to identify tumors that may respond to novel agents, including those that target tumors driven by *NF1* loss (Way, 2017). Although these machine learning algorithms are not ready for the clinic, the next logical step is to use PDX models to test the predictive nature of classifiers so that in the future, interdisciplinary teams can identify tumors driven by *TP53* and/or *NF1* loss, evaluate, and compare multiple therapies in real time.

Our study also highlights additional opportunities for pan-pediatric genomic characterization. We did not have available models for acute myelogenous leukemia, juvenile myelomonocytic leukemia, lymphomas, retinoblastoma, melanoma, or thyroid malignancies. Additionally, although we covered 29 histotypes, many of the rare tumors had low numbers and could benefit from creation and sequencing of additional models, and we seek to generate these data and/or hope to merge our data with future pediatric cancer PDX sequencing projects. Finally, WES likely missed several pathogenic lesions, and DNA methylation profiling is particularly relevant for pediatric brain tumors. Future studies, perhaps in collaboration with ongoing similar efforts by international colleagues, could address these gaps.

We performed this project to provide a resource to the pediatric cancer research community. To date, the pediatric cancer genomic literature largely focuses on diagnostic samples, and this study includes a large number of PDXs derived during or after intensive chemoradiotherapy. Thus, the frequency of many genomic alterations is higher in these models compared to the literature. By having a large number of PDXs obtained from samples at relapse or at autopsy, we can provide models that more closely recapitulate the patients being enrolled on early phase clinical trials after extensive chemoradiotherapy. All models are readily-available from the Children’s Oncology Group (www.CCCells.org) supported by ALSF or request to PI and data are openly-available from https://pedcbioportal.org/study?id=pptc for the pediatric oncology community.

## Supporting information

Rokita-PDX-Genomics-supplemental-tables

## Acknowledgments

The authors would like to thank the Alex’s Lemonade Stand Foundation for providing the funding for this project. This work was also supported by NIH grants U01 CA199287 (JMM), U01 CA199000 (RBL), U01 CA199288 (X-NL), U01 CA199221 (RG), U01 CA199297 (PJH), U01 CA199222 (GJG), R35 CA220500 (JMM), R01 CA221957 (CPR), NINDS R01 NS095411-01A1 (YS), the Giulio D’Angio Endowed Chair (JMM), the National Health and Medical Research Council of Australia (NHMRC Fellowships APP1059804 and APP1157871 to RBL and NHMRC Program Grant APP1091261), the Cancer Council New South Wales (PG 16-01), and Australian Federal Government Department of Health funding awarded to Zero Childhood Cancer, a joint initiative of Children’s Cancer Institute Australia (affiliated with UNSW Sydney) and The Kids Cancer Centre, Sydney Children’s Hospitals Network. We also acknowledge the NHLBI GO Exome Sequencing Project and its ongoing studies which produced and provided exome variant calls for comparison: the Lung GO Sequencing Project (HL-102923), the WHI Sequencing Project (HL-102924), the Broad GO Sequencing Project (HL-102925), the Seattle GO Sequencing Project (HL-102926) and the Heart GO Sequencing Project (HL-103010).

We acknowledge sequencing and molecular characterization efforts of the PPTP, with which our data was harmonized (Brian Geier, Dias Kurmashev). We also thank the following researchers for PDX model establishment and maintenance: Edward Favours, Doris Phelps, Alessia D’Aulerio, Colleen Larmour, and Matthew Tsang. The authors also thank Dr. David T. Teachey (Children’s Hospital of Philadelphia, Philadelphia, PA), Professor Charles G. Mullighan (St Jude Children’s Research Hospital, Memphis, TN) and the Children’s Oncology group for providing material from which the ETP-ALL and Ph-like ALL xenografts were established. We thank Chia Chin Wu, and Jianhua Zhang for helpful discussions throughout the study.

Finally, the authors gratefully acknowledge the patients and their families for donating samples for PDX establishment.

## Author Contributions

Conceptualization, JMM, DAW, JLR, MAS, PJH, RTK, X-NL, RBL, RG, MH

Methodology, DAW, JLR, JP, GPW, KSR

Software, KR, MFC, AF, JLR, GPW, KP, JP

Validation, CPR, JLR, KP, KLC, KAU, KM, JN, FKB

Formal Analysis, DAW, JLR, KSR, MFC, JP, LER, NMK, GPW, CM

Investigation, JLR, JP, KSR, KAU, KC, GPW, KLC, JN

Resources, JMGF, JB, KL, SC, ACB, JMM, KK, YPM, RBL, CSG, PJH, X-NL, RTK, HVD, ZM, JJ, KM, JN, CM, DK, KE, HM, JWB, KB, FKB, LQ, YD, HZ, HBL, SZ, JS, PB

Data curation, DAW, MC, JLR, KSR, KAU, KK, GIS, CM, KE, HM, KB, JWB

Writing – Original Draft, JLR, JMM, DAW, JJ, KSR, GPW, JP, NK, LER, KAU

Writing – Review & Editing, JLR, MAS, JMM, CSG, CM, RBL, YS, SZ, KK

Visualization, JLR, KSR, JP, NMK, KP, LER, GL, AM

Supervision, JMM, DAW, JLR, MAS, DH, CPR, SJD, OMV, ZV

Project Administration, JMM, JGF, JB, KL, MAS, GJG

Funding Acquisition, JMM, JGF, DAW, CPR, RBL, JLR, MH, GMM, VT, YS, RG, PJH, GJG

## CONTACT FOR REAGENT AND RESOURCE SHARING

Further information and requests for resources and reagents should be directed to and will be fulfilled by the Lead Contact, John M. Maris.

## EXPERIMENTAL MODEL AND SUBJECT DETAILS

### Patient-Derived xenograft generation and harvesting

Patient-derived xenograft models from the Pediatric Preclinical Testing Program (PPTP) were generated as described (Whiteford et al. 2007; Houghton et al. 2007; Houghton et al. 2002). Briefly, CB17/Icr scid-/-mice (Taconic Farms, Germantown NY), subcutaneously engraft kidney/rhabdoid tumors, sarcomas (Ewing, osteosarcoma, rhabdomyosarcoma), neuroblastoma, and non-glioblastoma brain tumors. For CNS tumors, patient tumor was surgically-transplanted into RAG2/SCID mouse brains in the diagnosis-specific orthotopic location as previously described (Yu et al. 2010). Whole murine brains containing visible tumors were aseptically removed and transferred to the tissue culture laboratory. Tumors were microscopically dissected from surrounding brain tissue, mechanically dissociated into cell suspensions, and filtered. Single tumor cells were subsequently injected into the brains of SCID mice as described above. Sub-transplantation process was repeated to complete a total of five tumor passages. All animal experiments were conducted according to an Institutional Animal Care and Use Committee-approved protocol. All leukemia animal experimentation was approved by the Animal Care and Ethics Committee, UNSW Sydney (Sydney, Australia). Experiments used continuous PDXs established previously in 20-25 g female non-obese diabetic/severe combined immuno-deficient (NOD.CB17-Prkdc^scid^/SzJ, NOD/SCID) or NOD/SCID/interleukin-2 receptor γ–negative (NOD.Cg-Prkdc^scid^ II2rg^tm^1^Wjl^/SzJ, NSG) mice, as described previously. Briefly, leukemia cells were inoculated intravenously into 6-8 week-old NOD/SCID or NSG mice (Australian BioResources, Moss Vale, NSW, Australia) and leukemia burden monitored via enumeration of human CD45^+^ (%huCD45^+^) cells versus total CD45^+^ leukocytes (human plus mouse) in the peripheral blood (PB) and tissues, as outlined previously (Lock et al. 2002; Liem et al. 2004). Additional details per model are included in Table S1.

## METHOD DETAILS

### Nucleic acid extractions and quality control

PDX samples were submitted from Children’s Cancer Institute, Children’s Hospital of Philadelphia, Greehey Children’s Cancer Research Institute, and Montefiore Medical Center to the Nationwide Children’s Hospital Biospecimen Core Resource at −190°C using an MVE cryoshipper. Cytospins and H&E frozen sections were prepared from leukemia and solid tissue PDX specimens, respectively. Slides were assessed by board-certified pathologists to determine blast percentage in leukemia PDX samples, and percent tumor nuclei and necrosis of the solid PDX samples. DNA and RNA were co-extracted from the PDXs using a modification of the DNA/RNA AllPrep kit (Qiagen). The flow-through from the Qiagen DNA column was processed using a mirVana miRNA Isolation Kit (Ambion). DNA was quantified by PicoGreen assay and RNA samples were quantified by measuring Abs_260_ with a UV spectrophotometer. DNA specimens were resolved by 1% agarose gel electrophoresis to confirm high molecular weight fragments. RNA was analyzed via the RNA6000 Nano assay (Agilent) for determination of an RNA Integrity Number (RIN). The PPTC study committee reviewed the pathology and molecular QC data and selected DNA and RNA aliquots for sequencing.

### Short tandem repeat (STR) profiling

Each tumor DNA sample was subjected to STR profiling performed by Guardian Forensic Sciences. DNA samples were quantified using Qiagen Investigator Quantiplex Kit (Cat# 387018) on a Qiagen RotorGene Q instrument. The GenePrint24 System for STR profiling (Promega, Cat#B1870) was used to amplify 0.05 ng of template DNA in a 12.5 µL volume using the following conditions: 96 °C for 1 minute, 27 cycles of {94 °C for 10 seconds, 59 °C for 1 minute, 72 °C for 30 seconds}, 60 °C for 10 minutes using the RotorGene Q instrument. Samples were injected into the Applied Biosystems ABI 310 Genetic Analyzer and profiles were interpreted by forensic biologists. Only those samples deemed not misidentified and free of contamination were used in this study.

### Biochemical measurement of human DNA content in PDX tumors

To determine the composition of human and mouse DNA within PDX tumors, PDX DNA samples were amplified using modified version of the published *pTGER2* (*prostaglandin E receptor 2*) qPCR assay (Alcoser et al. 2011). Depending upon sample availability, 2-20 ng of PDX tumor DNA were added to 500 nM each human- and mouse-specific forward primers, reverse primers, probes (sequences in resource document) and 1X IDT PrimeTime Gene Expression 2X Mastermix (Integrated DNA Technologies) in a total of 20 uL. Reactions were thermalcycled at 95 °C for 8 min and 42 cycles of {95 °C for 15 sec, 64 °C for 1 min}. Five-point standard curves were performed using a mixture of CHLA-90 and COG-N-603 neuroblastoma cell lines as human-specific template and pooled liver/spleen/muscle DNA from a naive NU/NU mouse as the mouse-specific template to confirm each primer efficiency was between 90-110%. The DNA equivalent of one diploid copy of either mouse or human template was run as a reference template. Three technical replicates were performed for each standard and sample. Average C_T_ values of the reference DNA samples were used as “ground truth” C_T_ values for one DNA copy.

To estimate relative copy number, 2^−ΔCT^.values were calculated for each unknown for each species:. To estimate percent human content, the following equation was used:.

### Whole exome sequencing

Illumina paired-end pre-capture libraries were constructed from PDX DNA samples according to the manufacturer’s protocol (Illumina Multiplexing_SamplePrep_Guide_1005361_D) modified as described in the BCM-HGSC Illumina Barcoded Paired-End Capture Library Preparation protocol. The complete protocol including oligonucleotide sequences used as adaptors and blockers are accessible from the HGSC website <https://www.hgsc.bcm.edu/sites/default/files/documents/Protocol->Illumina_Whole_Exome_Sequencing_Library_Preparation-KAPA_Version_BCM-HGSC_RD_03-20-2014.pdf. The DNA sequence production is described briefly below.

#### Library Preparation

500 ng (or 250 ng if sample quantity was limiting) of DNA in 50ul volume were sheared into fragments to an average size of 200-300 bp in a Covaris plate with E220 system (Covaris, Inc. Woburn, MA) followed by end-repair, A-tailing and ligation of the Illumina multiplexing PE adaptors. Pre-capture Ligation Mediated-PCR (LM-PCR) was performed for 6-8 cycles using the Library Amplification Readymix containing KAPA HiFi DNA Polymerase (Kapa Biosystems, Inc.). Universal primer LM-PCR Primer 1.0 and LM-PCR Primer 2.0 were used to amplify the ligated products. Reaction products were purified using 1.8X Agencourt AMPure XP beads (Beckman Coulter) after each enzymatic reaction. Following the final 1.2X Agentcourt XP beads purification, quantification and size distribution of the pre-capture LM-PCR product was determined using Fragment Analyzer capillary electrophoresis system (Advanced Analytical Technologies, Inc.).

#### Capture Enrichment

Four pre-capture libraries were pooled together (∼750 ng/sample, 3 ug/pool) and then hybridized in solution to the HGSC VCRome 2.1 design1 (Bainbridge et al. 2011) according to the manufacturer’s protocol NimbleGen SeqCap EZ Exome Library SR User’s Guide (Version 2.2) with minor revisions. Probes for exome coverage across >3,500 clinically relevant genes that are previously <20X (∼2.72Mb) is supplemented into the VCRome 2.1 probe. Human COT1 DNA was added into the hybridization to block repetitive genomic sequences. Blocking oligonucleotides from Sigma (individually sequence specifically synthesized) or xGen Universal Blocking oligonucleotides (Integrated DNA Technologies) were added into the hybridization to block the adaptor sequences. Hybridization was carried out at 560C for ∼16h. Post-capture LM-PCR amplification was performed using the Library Amplification Readymix containing KAPA HiFi DNA Polymerase (Kapa Biosystems, Inc.) with 12 cycles of amplification. After the final AMPure XP bead purification, quantity and size of the capture library was analyzed using the Agilent Bioanalyzer 2100 DNA Chip 7500. The efficiency of the capture was evaluated by performing a qPCR-based quality check on the four standard NimbleGen internal controls. Successful enrichment of the capture libraries was estimated to range from a 6 to 9 of ΔC_T_ value over the non-enriched samples.

#### DNA Sequencing

Library templates were prepared for sequencing using Illumina’s cBot cluster generation system with TruSeq PE Cluster Generation Kits (Illumina) according to the manufacturer’s protocol. Briefly, these libraries were denatured with sodium hydroxide and diluted to 6-9 pM in hybridization buffer in order to achieve a load density of ∼800K clusters/mm^2^. Each library pool was loaded in a single lane of a HiSeq flow cell, and each lane was spiked with 1% phiX control library for run quality control. The sample libraries then underwent bridge amplification to form clonal clusters, followed by hybridization with the sequencing primer. Sequencing runs were performed in paired-end mode using the Illumina HiSeq 2000 platform. Using the TruSeq SBS Kits (Illumina), sequencing-by-synthesis reactions were extended for 101 cycles from each end, with an additional 7 cycles for the index read. With sequencing yields averaging 12.1 Gb per sample, samples achieved an average of 97. 64% of the targeted exome bases covered to a depth of 20X or greater.

#### Primary Data Analysis

Initial sequence analysis was performed using the HGSC Mercury analysis pipeline (Challis et al. 2012; Reid et al. 2014). In summary, the .bcl files produced on-instrument were first transferred into the HGSC analysis infrastructure by the HiSeq Real-time Analysis module. Mercury then ran the vendor’s primary analysis software (CASAVA) to de-multiplex pooled samples and generate sequence reads and base-call confidence values (qualities), followed by the mapping of reads to the GRCh37 Human reference genome (http://www.ncbi.nlm.nih.gov/projects/genome/assembly/grc/human/) using the Burrows-Wheeler aligner (Li & Durbin 2010). The resulting BAM (binary alignment/map) file underwent quality recalibration using GATK, and where necessary the merging of separate sequence-event BAMs into a single sample-level BAM. BAM sorting, duplicate read marking, and realignment to improve in/del discovery all occur at this step. Next, Atlas-SNP and Atlas-indel from the Atlas2 suite (Shen et al. 2010) were used to call variants and produce a variant call file (VCF). Finally, annotation data was added to the VCF using a suite of annotation tools “Cassandra” (https://www.hgsc.bcm.edu/software/cassandra) that brings together frequency, function, and other relevant information using AnnoVar with UCSC and RefSeq gene models, as well as a host of other internal and external data resources.

### SNP array assay

In brief, 200 ng of genomic DNA were denatured with NaOH, followed by isothermal whole genome amplification at 37°C for 20-24 hours. The amplified DNA was enzymatically fragmented and hybridized to the BeadChip for 16-24 hours at 48°C (24 samples were processed in parallel for each BeadChip). After a series of washing steps to remove unhybridized and non-specifically hybridized DNA fragments, allele-specific single-base extension reactions were performed to incorporate labeled nucleotides into the bead-bound primers. A multi-layer staining process was conducted to amplify signals from the labeled extended primers, and then the coated beads were imaged with the Illumina iScan system.

Chip types used were humanomniexpress-24-v1-1-a.bpm and InfiniumOmniExpress-24v1-2_A1.bpm.

### Whole transcriptome sequencing

Whole-transcriptome RNA sequencing (RNA-seq) was performed using total RNA extracted as described above. Strand-specific, poly-A+ RNA-seq libraries for sequencing on the Illumina platform were prepared as previously described with minor modifications (Wang et al. 2015; Peters et al. 2015). RNA Integrity was confirmed (RIN >7.0) on a Bioanalyzer (Agilent). Briefly, poly-A+ mRNA was extracted from 1 μg total RNA using Oligo(dT)25 Dynabeads (Life Technologies), to which 4 µL of 1:100 dilution of the ERCC spike-in mix 1 (Ambion, Life technologies) was already added (Baker et al. 2005). There are a total of 92 poyladenylated transcripts in this mix that are used to monitor sample and process consistency. mRNA is then fragmented by heat at 94 °C for 15 minutes or less depending on sample RIN. First strand cDNA was synthesized using NEBNext RNA First Strand Synthesis Module (New England BioLabs) and during second strand cDNA synthesis, dNTP mix containing dUTP was used to introduce strand-specificity with NEBNext Ultra Directional RNA Second Strand Synthesis Module (New England BioLabs). For Illumina paired-end library construction, the resultant cDNA is processed through end-repair and A-tailing, ligated with Illumina PE adapters, and then digested with 10 units of Uracil-DNA Glycosylase (New England BioLabs). Libraries are prepared on the Beckman BioMek FXp robots and amplification of the libraries was performed for 13 PCR cycles using the Phusion High-Fidelity PCR Master Mix (New England BioLabs); 6-bp molecular barcodes that were also incorporated during this step. Libraries were purified with Agencourt AMPure XP beads (Beckman Coulter) after each enzymatic reaction, and after PCR amplification, and were quantified using Fragment Analyzer electrophoresis system. Libraries were pooled in equimolar amounts (4 libraries/pool). Library templates were prepared and sequenced exactly as described above for DNA Sequencing. Sequencing runs generated approximately 300-400 million successful reads on each lane of a flow cell, yielding 75-100M reads per sample.

## QUANTIFICATION AND STATISTICAL ANALYSIS

### Mouse read subtraction from WES sequencing data

Raw fastq files (N = 240) from Whole exome sequencing data were aligned to a combined hybrid genome of human hg19 and mouse mm10 genomes using the *Burrows-Wheeler transformation algorithm (BWA v0.7.17-r1188)*. Reads overlapping specifically to either the human or mouse genome were extracted and separated in corresponding human and mouse bam files using Samtools v1.9. The mouse subtracted bam files containing reads specific to human genome were then sorted by name and only paired reads were kept using the Samtools parameter *-f 1*. Following this, duplicated reads were removed using Sambamba v0.6.6. The deduplicated bam files were then used as input for local realignment around indels using IndelRealigner and base quality score recalibration using BaseRecalibrator utilities from GATK v3.8.1.

### Whole exome mutation analysis

Many of these PDX models have been established decades ago, thus matched primary and/or normal tissue either were not collected or is not currently available. To filter common germline variation from these tumor models, we used a panel of 809 normal samples supplied from TCGA WBC tissue to generate consensus germline variant calls. Rare germline variation was retained and defined as < 0.005 minor allele frequency in any one of the three databases: Exome Aggregation Consortium (ExAC) (Lek et al. 2016), 1000 genomes, or the NHBLI Exome Sequencing Project (ESP). Filtered variants also present in COSMIC were scavenged back. We performed MutSigCV (Lawrence et al. 2013) analysis on the entire cohort to identify and remove false positive variants. With the exception of known oncogenes and tumor suppressors, novel significantly mutated genes (SMGs) common across all histologies should be rare. We manually inspected the top 100 SMGs and found that most novel genes harbored a high number of private mutations and thus were not removed. Other novel variants were false positives due to germline inclusion or sequencing/mapping errors (Table S3). Data were thus split into germline MAF and somatic MAF files, the latter of which retained private variants.

### Literature mutation comparison analysis

The Pediatric Preclinical Testing Consortium (Maris, 2018) dataset in the PedCBio Portal was used to determine the gene alteration frequency in our cohort. For all studies, we tabulated only on non-silent single nucleotide variants since not all studies reported copy number alterations and fusions as separate entities. Previously reported gene alteration frequencies were calculated from the following literature studies. From Gröbner et al., “S_Table10” was used. We averaged sub-histologies “B-ALLOTHER” and “B-ALLHYPO” for comparison to PPTC PDX BCP-ALL. From Ma et al., the data in “TableS2” were used. The “Discovered by” column was filtered to only contained “GRIN & MutSIgCV” or “MutSigCV” and the value in the “Sample Counts (P/LP)” column was used to determine the proportion of tumors mutated. From Liu et al., “Table S9 Driver Mutations” was used. The column “#Sample” indicated the number of patients with a mutation for each gene and was used to determine the proportion. From Pugh et al., “Supplementary Table 3”, table entitled “Mutations” was used. Within this table, the “Hugo_Symbol” column was utilized; the repetitions for each gene were used to determine the proportion. From Eleveld et al., “Table S2” was filtered for relapsed samples, and the number of repetitions for each gene was used to determine the proportion. For all datasets, if a gene that was not included, it was deemed “NA” (Gröbner et al. 2018; Pugh et al. 2013; Y. Liu et al. 2017; Eleveld et al. 2015).

### Mutation correlation analyses

Plots correlating DNA variant allele frequencies for 13 diagnose-relapse model pairs and 6 model pairs belonging to the same phase were generated for several histologies and genes of interest (Table S5) were labelled. Total mutations by phase were visualized as stacked bar plots and boxplot.

### Indel *analysis*

Small structural variants were derived from Maftools (Mayakonda et al. 2018) (Supplemental Figure S1). Total number of frameshift and in-frame insertions and deletions (indels) were calculated per Model. Boxplots were generated with Histology on the x-axis and Number of indels per Model on y-axis.

### ATRX deletion analysis

The *ATRX* locus on chromosome X contains too few probes in OmniExpress arrays to accurately assess deletion, even in cases of known sex. Thus, from WES bam files, total read base counts for ATRX exons were calculated using Samtools v1.9 bedcov utility and total library size was calculated using Samtools v1.9 flagstat utility. To convert exon read counts to Fragments per kilobase per million reads (FPKM), the library sizes were first transformed to per million scaling factors. Following this, raw read counts of each exon were normalized using the per million scaling factors and the corresponding exon length.

### Mutational signatures analysis

The deconstructSigs R package with the COSMIC 30 signature reference was used. We plotted only models with ≥50 total somatic mutations. We quantified the proportion of models within each histology having signatures with a cosine similarity value > 0.1. These proportions were grouped into one of four cosmic signature categories based on aetiology (5-mC deamination, APOBEC, DNA repair, Hypermutation and/or MSI) and plotted as stacked bar graphs.

### Classifier analysis

We applied models derived from three supervised machine learning algorithms to all PDX models with available RNA-Seq data (N = 244). The models were previously trained on RNAseq, copy number, and mutation data across 33 different adult cancer-types from The Cancer Genome Atlas PanCanAtlas project (Cancer Genome Atlas Research Network et al. 2013). Briefly, the algorithm was an elastic net penalized logistic regression classifier that took FPKM and z-score normalized RNAseq data as input and, in three independent classifiers, was trained to predict Ras pathway activation, *NF1* inactivation, and *TP53* inactivation using mutation and copy number alteration status of corresponding samples. The Ras pathway and *NF1* classifiers and the overall method were described in more detail in (Way et al. 2018). The application and validation of the *TP53* classifier was described in (Knijnenburg et al. 2018).

To assess performance of the TCGA trained classifiers applied to the PDX data, we used orthogonal evidence of gene alterations in each PDX sample. Specifically, we used samples with observed missense, nonsense, frame shift, and splice site mutations in *ALK, BRAF, CIC, DMD, HRAS, KRAS, NF1, NRAS, PTPN11*, and *SOS1* as samples with possible Ras pathway activation. We used samples with only non-silent *NF1* mutations for the *NF1* classifier, and samples with deleterious *TP53* mutations, copy number deletions, and fusions for the *TP53* classifier. We assessed model performance using receiver operating characteristic (ROC) and precision recall (PR) curves using these samples as the positive set and all others as the negative set. We also applied the classifiers to shuffled PDX gene expression matrices and compared performance to the real data to assess potential model bias. The reproducible analysis pipeline can be viewed at https://github.com/marislab/pdx-classification and the software is archived on Zenodo at doi: 10.5281/Zenodo.1475249.

### mRNA gene expression analysis

Raw fastq files (N = 244) from RNA-sequencing data were aligned to a combined hybrid genome of human hg19 and mouse mm10 genomes using the STAR aligner v2.5.3a. Reads overlapping specifically to either the human or mouse genome were extracted and separated in corresponding human and mouse bam files using Samtools v1.9. The mouse subtracted bam files containing reads specific to human genome were then sorted by name and only paired reads were kept using the Samtools parameter *-f 1*. Following this, duplicated reads were removed using Sambamba v0.6.6. The deduplicated bam files were used to extract and separate reads into paired-ended fastq files using the *SamToFastq* utility of Picard v2.18.14-0. The resulting paired-ended fastq files obtained after mouse subtraction were re-aligned to human genome hg19 using STAR aligner and filtered for duplicate reads using Picard *MarkDuplicates*. Gene expression was quantified in terms of Fragments Per Kilobase of transcript per Million mapped reads (FPKM) using HTSeq v0.9.1 and Cufflinks v2.2.1. We also processed RNA-sequencing patient data from TARGET (ALL, N = 533; AML, N = 364; NBL, N = 169; RT, N = 70; OS, N = 87; WT, N = 136) and PPTC PDX data (N = 244) using STAR alignment and RSEM normalization using hg38 as reference genome and Gencode v23 gene annotation to get transcript per million (TPM) expression values. For PPTC PDX data, human bam files generated from the mouse subtraction pipeline were used in order to generate input fastq files.

### mRNA variant calling, filtering, and comparison to DNA variants

Variant calling for RNA-seq samples was performed with Strelka v2.9.2 germline indels calling pipeline using hg19 primary assembly reference fasta and default parameters. VCFs were converted to MAF and variants were filtered for those that passed VEP and were non-silent (!= Silent or Intron). Variant allele frequencies for all non-silent, VEP-passed RNA variants were calculated. For each model on which both WES and RNA-Seq were performed, WES variants with RNA evidence were matched in the DNA MAF and VAF correlations were plotted.

### Copy number analysis

SNP arrays were processed at the HGSC using the Illumina Infinium HTS Assay according to the manufacturer’s guidelines. Human OmniExpress arrays (Illumina, catalog No. WG-315-1101) were used, interrogating 741 thousand SNP loci with a MAF detection limit of 5%. SNP calls were collected using Illumina’s GenomeStudio software (version 1.0/2.0) in which standard SNP clustering and genotyping were performed with the default settings recommended by the manufacturer. Data from samples that met a minimum SNP call rate of 0.9 were considered passing and were included in subsequent analyses. Output files from Genome Studio containing BAF and LRR were used as input for Nexus 8.0. Quadratic systematic correction was performed using a custom file (Figshare repository, below) containing common snp probes from the two chip types. The significance threshold was reduced to 1 x 10^−8^ to reduce background noise. Segmentation was performed using Nexus’s SNPRANK algorithm. To extract segments, gain was set to 0 and loss to −1 x 10^−11^. The output table was reformatted to segmentation file format for input to GISTIC2.0, which was used to calculate broad and focal, hemizygous gene-level copy number events. Relevant arm and band level alterations were used in oncoprints. Since normal DNA was not available for paired analyses, sex chromosomes were removed. Focal homozygous deletions and amplifications were annotated using the segmentation file created post-Nexus analysis. A cutoff of LRR >=(0.538578182) was used for amplifications and >=(−1.739) for deletions. Cutoffs were determined by assessing histogram splits for MYCN amplification, SMARCB1 deletion, and CDKN2A/B deletions. Homozygous deletions remained only if mRNA FPKM was < 5 or if RNA-Seq for a sample was not available. Manual inspections were performed to confirm alterations for *SMARCB1, TP53, WT1, MYCN, C19MC, CDKN2A/B* and edited when necessary (see code).

### Ethnicity inference

Approximate genomic ancestries for each PDX model were inferred through principal component analysis of SNP array genotypes. Illumina-designated plus-strand genotypes were exported from GenomeStudio and processed using PLINK 1.9. Sex chromosomes and SNPs with minor allele frequency <1%, call rate <90%, or a deviation from Hardy-Weinberg equilibrium surpassing p=0.00005 were excluded. The PDX dataset was then merged with HapMap 3 (draft release 2), restricting to only the intersecting SNPs. This set was pruned to remove highly correlated SNPs using a window size of 50 variants, step size of 5 variants, and pairwise r^2^ threshold of 0.1. The 39,544 remaining SNPs were used to calculate the top 20 principal components. Approximate ethnicities were inferred using the first two components. Individuals were classified into four broad population groups: European (including HapMap CEU and TSI population samples), African (ASW, LWK, MKK, and YRI), East Asian (CHB, CHD, and JPT), and South Asian or Hispanic (GIH and MXL).

### Fusion transcript analysis

We used four different fusion callers: STAR-Fusion v1.1.0, FusionCatcher v0.99.7b, deFuse and SOAPFuse on RNA-sequencing data of the PDX models (N = 244). A total of 50,796 unique fusions were predicted with the following breakdown: STAR-Fusion (N = 9,496), FusionCatcher (N = 3,822), deFuse (N = 30,393), and SOAPFuse (N = 7,085). To reduce the number of false positives, we used two parallel approaches: first to keep all fusions predicted as in-frame and second to keep all fusions where the 5’ or 3’ gene fuses promiscuously with multiple partners within the same histology. To filter out unreliable predictions, we further filtered the in-frame fusions by keeping fusions that were recurrently predicted in two or more models within the sample histology or fusions that were supported by at least two fusion callers. We removed any fusions where expression of both genes in the gene pair was found to be < 1 TPM value across all models or it was not reported by the gene quantification algorithm. We then combined the lists from the two approaches discussed above and filtered out any fusions that were predicted in more than one histology. To remove spurious fusions, we filtered all fusions annotated as “read-through” as a result of fusions between adjacent or neighboring genes. We further removed fusions identified in non-cancer tissues and cells as per GTEx in order to remove chimeric RNA that is normally found in healthy tissue. Next, we scavenged and annotated fusions that have been identified as “driver” fusions in literature and fusions that were validated using cytogenetics. Finally, we annotated the gene fusion partners with oncogenes from COSMIC, kinases from Kinase.com, and transcription factors from AnimalTFDB to identify any oncogenic potential and functional relevance.

### RNA expression clustering and pathway analyses

The UCSC TumorMap analysis was used to visualize clusters of expression profiles across PDX histologies (Newton et al. 2017). The expression values were transformed into log2(TPM + 1) space. We removed genes where more than 80% of the samples had no measurable expression and we applied a variance filter to remove the 20% least varying genes. This generated a gene by sample matrix containing 28,482 genes and 244 PDX samples. The expression values and PDX annotations were uploaded to the TumorMap portal for analysis. A Bayesian hierarchical model was used to infer differences in expression across PDX histologies. We used a hierarchical modeling strategy to leverage similarities across related tissues and to improve inferences for histologies with small sample sizes (Ji & X. S. Liu 2010). The hierarchical model was implemented using the Stan statistical programming language (Carpenter et al. 2017).

We inferred the biological function of histology-specific expression by ranking the expression differences for each histology and performing gene-set enrichment analysis (GSEA). GSEA was performed using the fgsea software {BioRxiv:ge}. Statistically significant enrichment was defined as having an adjusted p-value less than 0.01 and a normalized enrichment score greater than 2.0. Statistically insignificant enrichment scores were set to zero for heatmap visualization. The normalized enrichment scores were visualized using the seaborn clustermap software (Waskom n.d.) for tissue database scores and R for Hallmark pathway scores.

### Pediatric cBioPortal data processing

All processed data: RNA-sequencing expression values (FPKM and Z-score), RNA fusions, mutation calls in Mutation Annotation Format (MAF), segmentation, and focal copy number values were formatted using the current cBioPortal v1.2.2 file format documentation (https://cbioportal.readthedocs.io/en/latest/File-Formats.html)

## DATA AND SOFTWARE AVAILABILITY

### Raw data availability

Mouse and human separated DNA and RNA BAM files have been deposited into dbGAP under accession number phs001437.

### Intermediate processed data availability

Variant files, SNP array files, contamination assessment files: https://figshare.com/projects/Genomic_landscape_of_childhood_cancer_patient-derived_xenograft_models/38147

Classifier Figshare files: https://figshare.com/articles/PPTC_RNAseq_and_Genomic_Alterations/7127726

### Processed data availability

WES mutations, mRNA expression, RNA fusions, segmentation, and gene copy number has been deposited into the publicly-available pediatric cBioportal at: https://pedcbioportal.org/study?id=pptc#summary

### Code created or modified for analysis in this paper have been deposited in GitHub

PDX mouse subtraction: https://github.com/marislab/pdx-mouse-subtraction

Ethnicity inference: https://github.com/lauraritenour/pptc-pdx-ethnicity-inference

Oncoprint generation: https://github.com/marislab/create-pptc-pdx-oncoprints

Gene classification: https://github.com/marislab/pdx-classification

RNA clustering and heatmaps: https://github.com/marislab/pptc-pdx-RNA-Seq-clustering

RNA fusion analysis: https://github.com/marislab/pptx-pdx-fusion-analysis

Copy number and SV analysis: https://github.com/marislab/pptc-pdx-copy-number-and-SVs

Correlation analyses: https://github.com/marislab/create-pptc-pdx-corplots

Mutational signatures: https://github.com/marislab/pptc-pdx-mut-sigs

## Supplemental Figure and Table Legends

**Figure S1.**
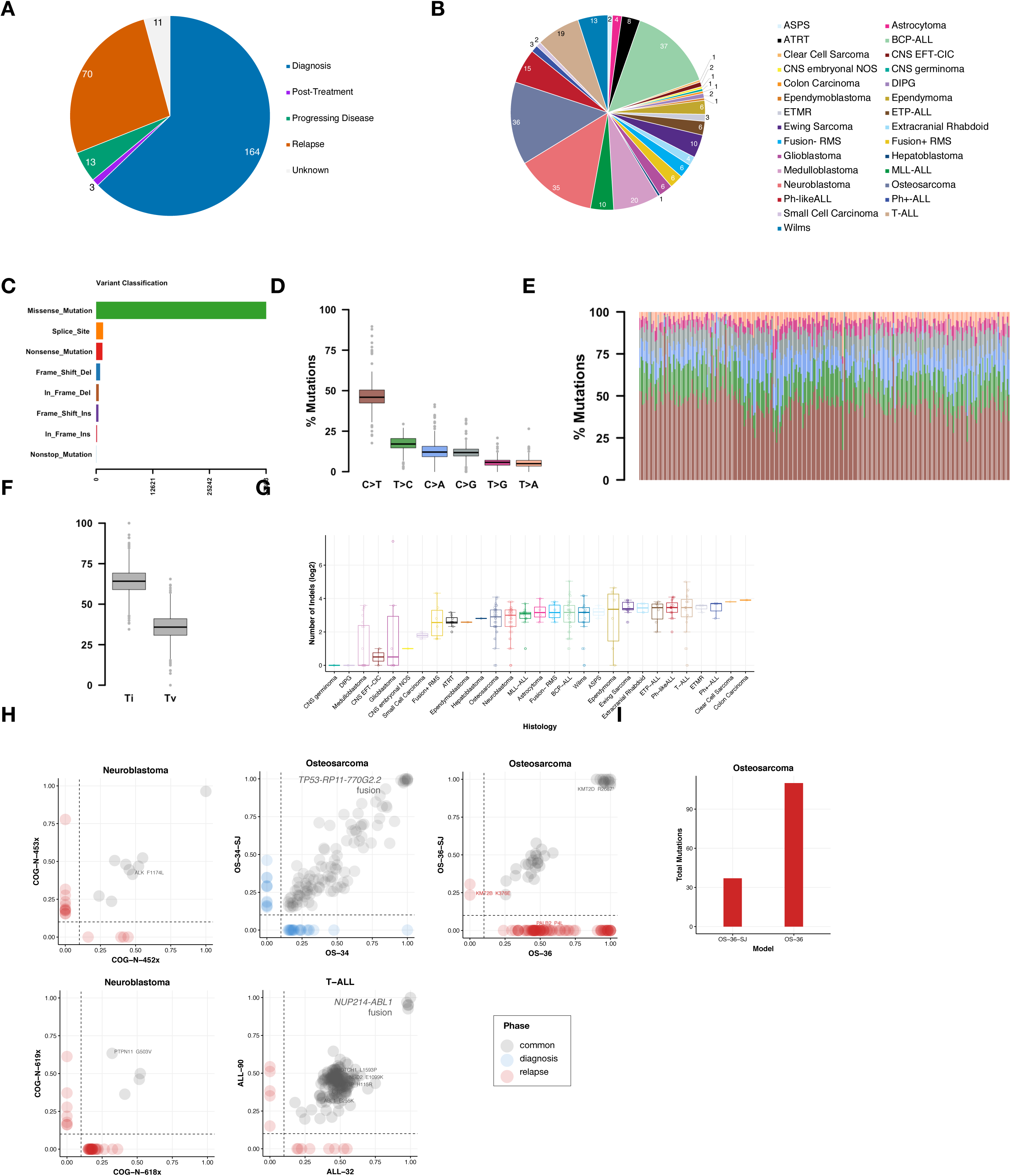
Histology and mutational breakdown, Related to Figure 1>. Pie charts showing breakdown of phase of therapy (A) and histology (B) for 261 PDX models. Variant classification breakdown (C), variant type breakdown (D), percent of mutations by nucleotide change (E), percent transitions (Ti) and transversions (Tv) (F), histogram of variants per sample (median = 133) (G), and top 10 genes mutated (H, x-axis = number of mutations, colors explained by C). Median number of small insertions and deletions (indels) were plotted per histology (I, box edges as first and third quartiles; detailed Ns in Table S4). J, same-phase scatterplots for model pairs created at the same phase of therapy (blue = diagnostic samples, red = relapse samples, grey = common mutations between two models plotted), K, total mutations in OS-26 and OS-36-SJ relapse samples.

**Figure S2.**
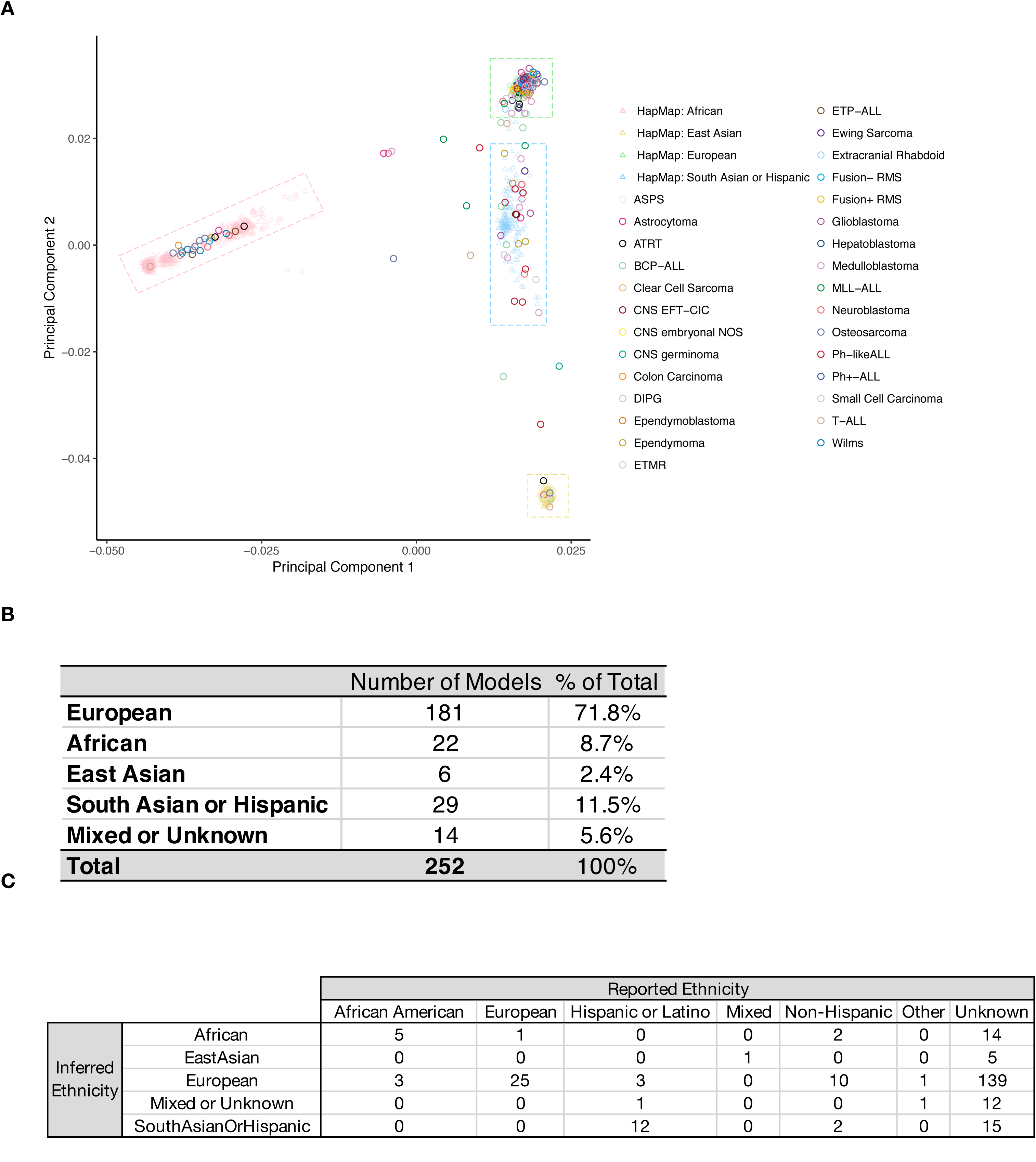
Ethnicity prediction, Related to Figure 1. Principal components analysis grouping of European, African, East Asian, and South Asian/Hispanic HapMap reference populations used to predict PDX ethnicities (A). The first two principal components calculated from SNP array genotypes for 252 PDX models (circles) are plotted alongside 1,184 HapMap reference samples (triangles). Dashed boxes represent the cutoffs used to classify PDXs into four broad population groups: European (including HapMap CEU and TSI population samples), African (ASW, LWK, MKK, and YRI), East Asian (CHB, CHD, and JPT), and South Asian or Hispanic (GIH and MXL). Tabulated counts and frequencies of ethnicities in PDX cohort (B) and a comparison table of reported versus inferred ethnicities in the PDX cohort (C).

**Figure S3.**
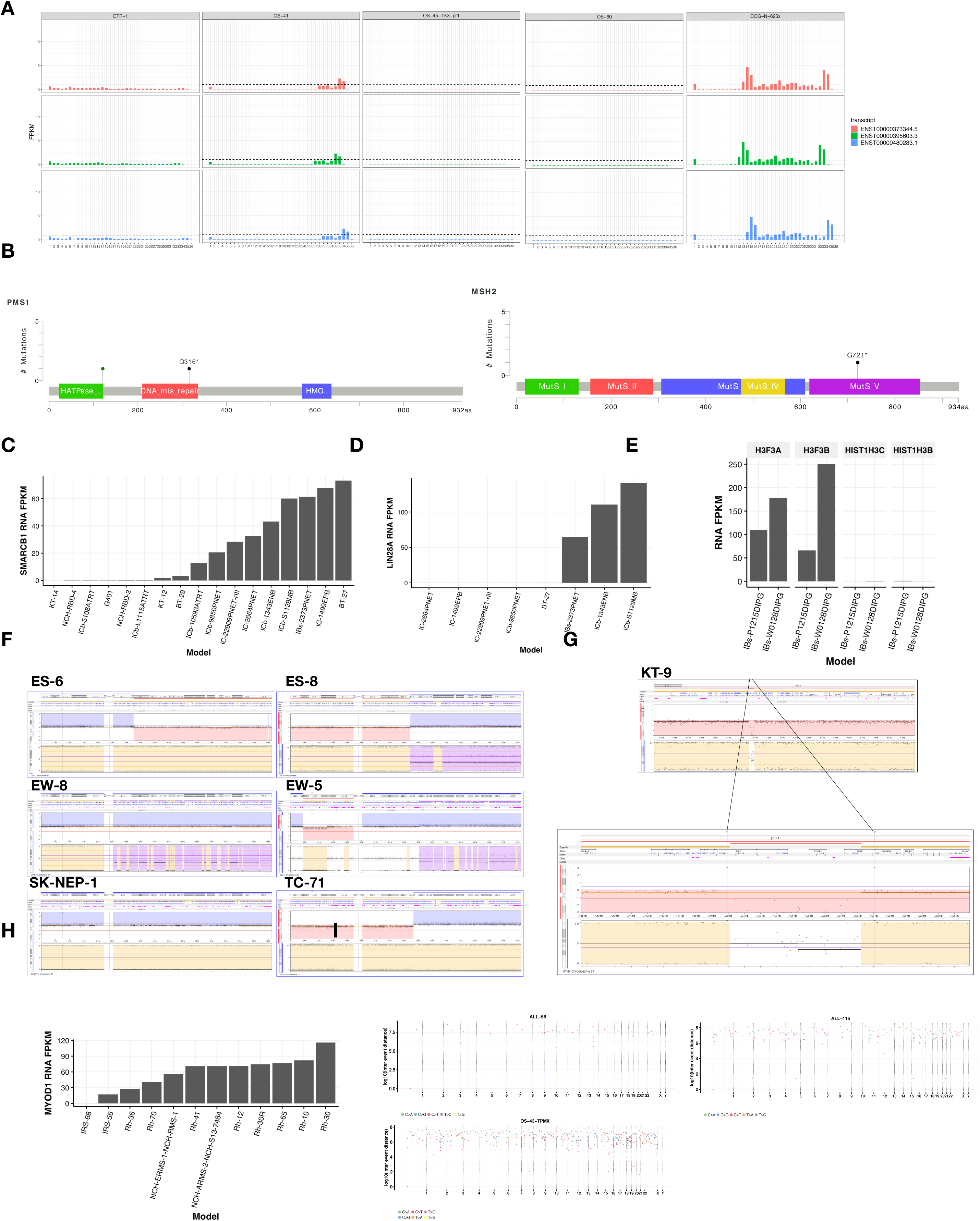
Genetic alteration evidence, Related to Figure 2. *ATRX* expression by exon for transcripts ENST00000373344.5, ENST00000395603.3, and ENST0000480283.1 derived from WES showing deletions in ETP-1, OS-41, OS-45-TSX-pr1, OS-60, and COG-N-625x (A, STAR Methods). Lollipop plots for ocogenic mutations in DNA repair genes, *PMS1* and *MSH2* for hypermutated model, IC-1621GBM (B). Gene expression for *SMARCB1* across ATRT and previously-classified PNET models (C), *LIN28A* for previously-classified PNET models (D), and *H3F3A, H3F3B, HIST1H3B,* and *HIST1H3C* for DIPG models (E). Nexus screenshots of the *TP53* locus in Ewing’s tumor models to validate homozygous or hemizygous deletions and loss of heterozygosity (F). Nexus screenshot of *TP53* deletion and breakpoints for *TP53-FXR2* fusion in KT-9 (G). *MYOD1* gene expression across rhabdomyosarcoma models (H). Rainfall plots for samples with predicted APOBEC signatures: ALL-58, ALL-115, and OS-43-TPMX (I).

**Figure S4.**
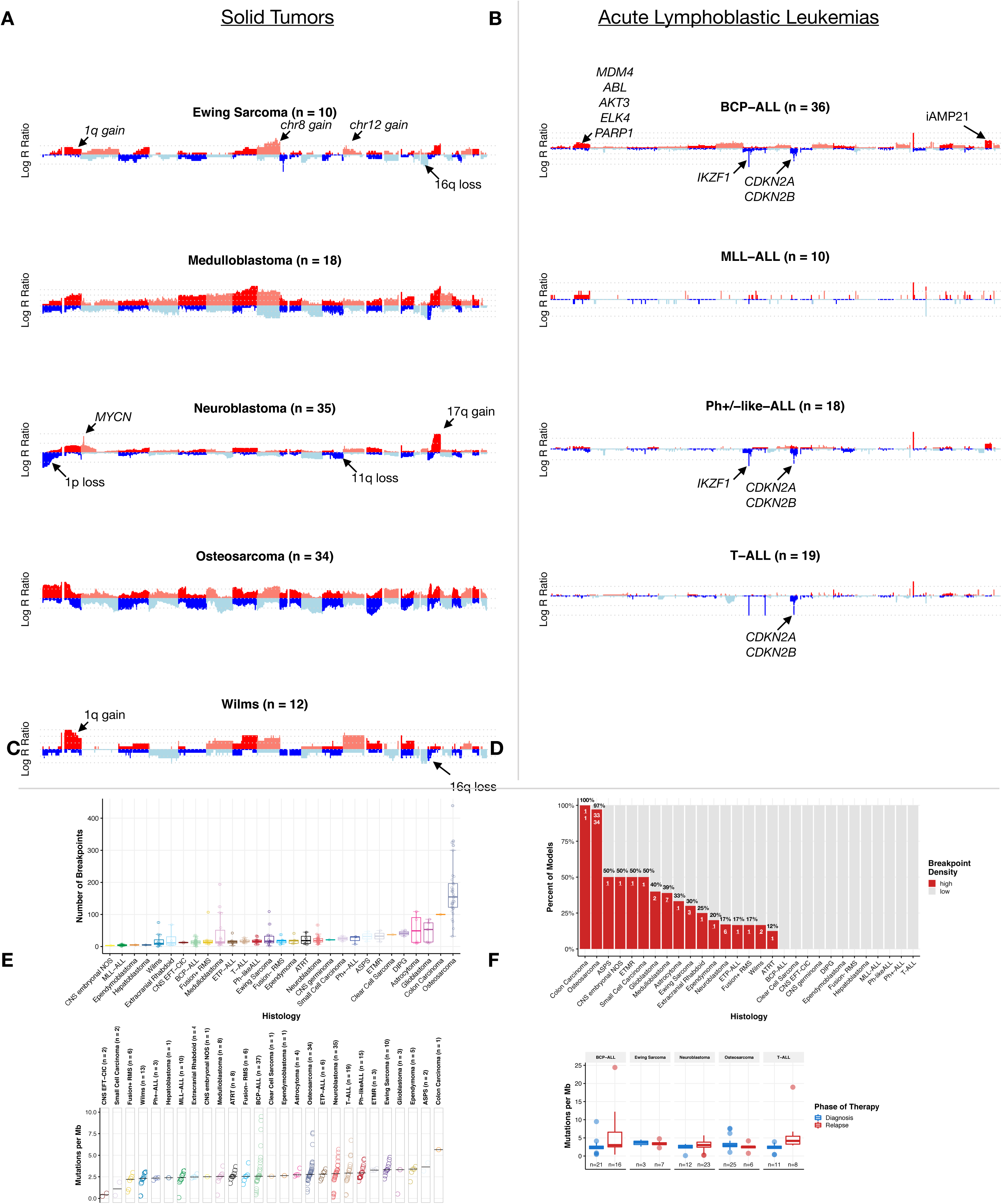
Copy number, TMB, and tumor evolution. Related to Figures 2 and 3. Figure 5 depicts genome-wide copy-number profiles for histologies with N ≧ 10 models (Panel A, solid tumors: Ewing sarcoma, N = 10; Medulloblastoma, N = 18; Neuroblastoma, N = 35; Osteosarcoma, N = 34; Wilms, N = 12 and Panel B, leukemias: BCP-ALL, N = 36; MLL-ALL, N = 10; Ph+ or Ph-like ALL, N = 18; T-ALL, N = 19). Canonical broad and focal lesions are annotated by histology. Breakpoints per histology are plotted in C (boxplots are graphed as medians with box edges as first and third quartiles; detailed Ns in Table S4) and breakpoint density across histologies is plotted in D (displayed as percent of models per histology and N / total; details in Table S4). Tumor mutation burden by histology across 240 models on which WES was performed (E, STAR methods). Histologies are plotted in rank order by median (y-intercept) and Ns per histology are listed. Tumor mutation burden by phase of therapy for histologies containing models from diagnosis and relapse (F). Boxplots represent median and first and third quartiles; Ns listed.

**Figure S5.**
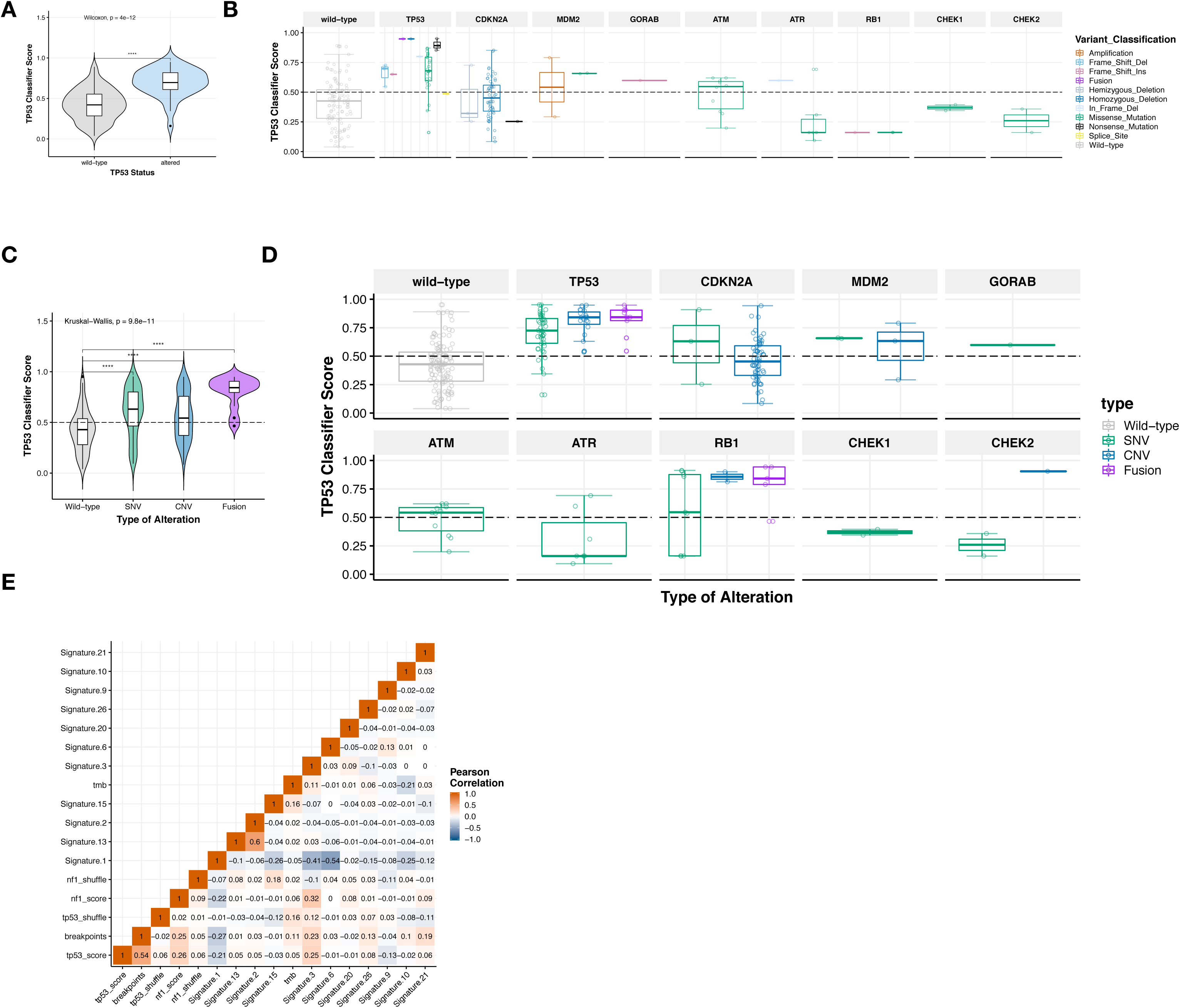
Classifier scores and mutational signature correlations, Related to Figure 4. With osteosarcoma models removed from analysis, *TP53* classifier scores were still significantly higher (Wilcoxon p = 4e-12) in models with a *TP53* alteration (A), but alterations in other pathway genes don’t consistently phenocopy *TP53* inactivation (B). Models containing fusions had highest classifier scores, followed by models with SNVs and CNVs, respectively (C, Kruskal-Wallis p = 9.8e-11) and these are broken down by gene in panel D. Validation of mutational signatures via Pearson correlation matrix: Signatures 2 and 13 correlate strongly (R = 0.6), Signature 1 is inversely correlated with impaired DNA repair mutational signatures, 3 (R = −0.41) and 6 (R = −0.54) (E).

**Table S1. Clinical, mouse, and metadata associated with 261 PDX models. Related to Figures 1, 2, and 3.** Breakdown of models per histology by assay performed and tables of clinical, mouse, and data file information for the PDX models described in this paper. Paired samples derived from the same patient are listed.

**Table S2. STR profiles for 261 PDX models. Related to Figure 1.** Geneprint 24 short tandem repeat profiles for all models described in this paper, along with all available laboratory, web, and published reference profiles.

**Table S3. Genes inspected for artifacts. Related to Figure 2.** MutSigCV results and determination of artifactual mutations in genes were annotated.

**Table S4. Summary mutation and demographics per model. Related to Figures 2, 5, and 6.** Mutations per model, mutations per megabase, breakpoints, high breakpoint density, and fusions are listed per model. Ages are summarized by histology. Summary statistics are listed within the table.

**Table S5. Driver gene of interest lists, driver fusions, and oncoprint matrices. Related to Figure 2.** Lists of driver genes and literature sources used for oncoprint plots. The rare histology oncoprint was created using all genes as input. Genes creating canonical oncogenic fusions or known oncogenic fusions are listed per histology. Oncoprint matrices for Figure 2 are provided.

**Table S6. Classifier scores for *TP53, NF1,* and Ras. Related to Figure 4.** Listed are all osteosarcoma tumors on which RNA-Seq was performed (N = 30). All except OS-55-SBX (score = 0.44) had scores predictive of *TP53* pathway inactivation. Genetic alterations within *TP53, CDKN2A, MDM2, MDM4, GORAB, ATM, ATR, RB1, CHEK1,* and *CHEK2* are listed as potential evidence of *TP53* inactivation in these models. Classifier and scores resulting from a shuffled gene expression matrix for each PDX model for which RNA-Seq was performed (N = 244). Raw, unfiltered COSMIC 30 mutational signature weights for all PDX models for which WES was performed (N = 240).

**Table S7. High-confidence fusion predictions. Related to Figure 5.** Filtered driver gene fusions and high-confidence fusions for which at least one gene partner was expressed (pipeline in Figure 1D and STAR methods). Fusions were annotated for being a kinase, transcription factor, or gene with oncogenic potential. FPKM of each fusion partner is listed and fusions without any expression were removed.

## References

Alexandrov, L.B. et al., 2013. Signatures of mutational processes in human cancer. Nature, 500(7463), pp.415–421.

American Childhood Cancer Organization, 2014. Special Section: Cancer in Children & Adolescents,

Behjati, S. et al., 2017. Recurrent mutation of IGF signalling genes and distinct patterns of genomic rearrangement in osteosarcoma. Nature Communications, 8, p.15936.

Boeva, V. et al., 2013. Breakpoint features of genomic rearrangements in neuroblastoma with unbalanced translocations and chromothripsis. PloS one, 8(8), p.e72182.

Bosse, K.R. et al., 2017. Identification of GPC2 as an Oncoprotein and Candidate Immunotherapeutic Target in High-Risk Neuroblastoma. Cancer cell, 32(3), pp.295–309.e12.

Cortes-Ciriano, I. et al., Comprehensive analysis of chromothripsis in 2,658 human cancers using whole-genome sequencing. biorxiv.org

El-Hoss, J. et al., 2016. A single nucleotide polymorphism genotyping platform for the authentication of patient derived xenografts. Oncotarget, 7(37), pp.60475–60490.

Eleveld, T.F. et al., 2015. Relapsed neuroblastomas show frequent RAS-MAPK pathway mutations. Nature Genetics, 47(8), pp.864–871.

Gelman, A., 2006. Multilevel (hierarchical) modeling: what it can and cannot do. Technometrics, 48(3), pp.432–435.

Gröbner, S.N. et al., 2018. The landscape of genomic alterations across childhood cancers. Nature, pp.1–23.

Houghton, p.J. et al., 2002. Testing of new agents in childhood cancer preclinical models: meeting summary. Clinical Cancer Research, 8(12), pp.3646–3657.

Houghton, P.J. et al., 2007. The pediatric preclinical testing program: description of models and early testing results. Pediatric Blood & Cancer, 49(7), pp.928–940.

Kim, P., Park, A., Han, G., Sun, H., Jia, P. and Zhao, Z., 2017. TissGDB: tissue-specific gene database in cancer. Nucleic acids research, 46(D1), pp.D1031–D1038.

Knijnenburg, T.A. et al., 2018. Genomic and Molecular Landscape of DNA Damage Repair Deficiency across The Cancer Genome Atlas. CellReports, 23(1), pp.239–254.e6.

Lecona, E. & Fernandez-Capetillo, O., 2018. Targeting ATR in cancer. Nature Reviews Cancer, 18(9), pp.586–595.

Liu, X., Yu, X., Zack, D.J., Zhu, H. and Qian, J., 2008. TiGER: a database for tissue-specific gene expression and regulation. BMC bioinformatics, 9(1), p.271.

Liu, Y. et al., 2017. The genomic landscape of pediatric and young adult T-lineage acute lymphoblastic leukemia. Nature Genetics, 49(8), pp.1211–1218.

Lock, R.B. et al., 2018. Abstract LB-B16: The AKR1C3-Activated Prodrug OBI-3424 Exerts Profound In Vivo Efficacy Against Preclinical Models of T-Cell Acute Lymphoblastic Leukemia (T-ALL); a Pediatric Preclinical Testing Consortium Study.

Lorenz, S. et al., 2016. Unscrambling the genomic chaos of osteosarcoma reveals extensive transcript fusion, recurrent rearrangements and frequent novel TP53 aberrations. Oncotarget, 7(5), pp.5273–5288.

Ma, X. et al., 2018. Pan-cancer genome and transcriptome analyses of 1,699 paediatric leukaemias and solid tumours. Nature, 555(7696), pp.371–376.

Ma, X. et al., 2015. Rise and fall of subclones from diagnosis to relapse in pediatric B-acute lymphoblastic leukaemia. Nature Communications, 6, p.6604.

Molenaar, J.J. et al., 2012. Sequencing of neuroblastoma identifies chromothripsis and defects in neuritogenesis genes. Nature, 483(7391), pp.589–593.

Newton, Y. et al., 2017. TumorMap: Exploring the Molecular Similarities of Cancer Samples in an Interactive Portal. Cancer Research, 77(21), pp.e111–e114.

Northcott, P.A. et al., 2017. The whole-genome landscape of medulloblastoma subtypes. Nature, 547(7663), pp.311–317.

O’Sullivan, C.G. et al., 2018. A Phase II Study of Atezolizumab in Patients with Alveolar Soft Part Sarcoma. In Connective Tissue Oncology Society Annual Meeting.

Padovan-Merhar, O.M. et al., 2016. Enrichment of Targetable Mutations in the Relapsed Neuroblastoma Genome. PLoS Genetics, 12(12), p.e1006501.

Pan, Z. et al., 2017. Loss of heterozygosity on chromosome 16q increases relapse risk in Wilms’ tumor: a meta-analysis. Oncotarget, 8(39), pp.66467–66475.

Pugh, T.J. et al., 2013. The genetic landscape of high-risk neuroblastoma. Nature Genetics, 45(3), pp.279–284.

Rausch, T. et al., 2012. Genome sequencing of pediatric medulloblastoma links catastrophic DNA rearrangements with TP53 mutations. Cell, 148(1-2), pp.59–71.

Ribi, S. et al., 2015. TP53 intron 1 hotspot rearrangements are specific to sporadic osteosarcoma and can cause Li-Fraumeni syndrome. Oncotarget, 6(10), pp.7727–7740.

Sano, R. et al., 2018. Abstract LB-136: Pediatric Preclinical Testing Consortium evaluation of a DLL3-targeted antibody drug conjugate rovalpituzumab tesirine, in neuroblastoma.

Schleiermacher, G. et al., 2014. Emergence of new ALK mutations at relapse of neuroblastoma. Journal of Clinical Oncology, 32(25), pp.2727–2734.

Schramm, A. et al., 2015. Mutational dynamics between primary and relapse neuroblastomas. Nature Genetics, 47(8), pp.872–877.

Scott, R.H. et al., 2012. Stratification of Wilms tumor by genetic and epigenetic analysis. Oncotarget, 3(3), pp.327–335.

Segers, H. et al., 2013. Gain of 1q is a marker of poor prognosis in Wilms’ tumors. Genes, Chromosomes and Cancer, 52(11), pp.1065–1074.

Shern, J.F. et al., 2014. Comprehensive genomic analysis of rhabdomyosarcoma reveals a landscape of alterations affecting a common genetic axis in fusion-positive and fusion-negative tumors. Cancer Discovery, 4(2), pp.216–231.

Spreafico, F. et al., 2013. Loss of heterozygosity analysis at different chromosome regions in Wilms tumor confirms 1p allelic loss as a marker of worse prognosis: a study from the Italian Association of Pediatric Hematology and Oncology. The Journal of urology, 189(1), pp.260–266.

Tirode, F. et al., 2014. Genomic landscape of Ewing sarcoma defines an aggressive subtype with co-association of STAG2 and TP53 mutations. Cancer Discovery, 4(11), pp.1342– 1353.

Way, G.P. et al., 2017. A machine learning classifier trained on cancer transcriptomes detects NF1 inactivation signal in glioblastoma. BMC Genomics, 18(1), p.127.

Way, G.P. et al., 2018. Machine Learning Detects Pan-cancer Ras Pathway Activation in The Cancer Genome Atlas. CellReports, 23(1), pp.172–180.e3.

Whiteford, C.C. et al., 2007. Credentialing preclinical pediatric xenograft models using gene expression and tissue microarray analysis. Cancer Research, 67(1), pp.32–40.

Zhang, J. et al., 2012. The genetic basis of early T-cell precursor acute lymphoblastic leukaemia. Nature Reviews Disease Primers, 481(7380), pp.157–163.

## METHOD REFERENCES

Alcoser, S.Y. et al., 2011. Real-time PCR-based assay to quantify the relative amount of human and mouse tissue present in tumor xenografts. BMC biotechnology, 11(1), p.124.

Bainbridge, M.N. et al., 2011. Targeted enrichment beyond the consensus coding DNA sequence exome reveals exons with higher variant densities. Genome Biology, 12(7), p.R68.

Baker, S.C. et al., 2005. The External RNA Controls Consortium: a progress report. Nature Methods, 2(10), pp.731–734.

Cancer Genome Atlas Research Network et al., 2013. The Cancer Genome Atlas Pan-Cancer analysis project. Nature Genetics, 45(10), pp.1113–1120.

Carpenter, B. et al., 2017. Stan: A probabilistic programming language. Journal of Statistical Software, 76(1), pp.1–32.

Challis, D. et al., 2012. An integrative variant analysis suite for whole exome next-generation sequencing data. BMC Bioinformatics, 13(1), p.8.

Houghton, P.J. et al., 2002. Testing of new agents in childhood cancer preclinical models: meeting summary. Clinical Cancer Research, 8(12), pp.3646–3657.

Ji, H. & Liu, X.S., 2010. Analyzing ’omics data using hierarchical models. Nature Biotechnology, 28(4), pp.337–340.

Lawrence, M.S. et al., 2013. Mutational heterogeneity in cancer and the search for new cancer-associated genes. Nature, 499(7457), pp.214–218.

Lek, M. et al., 2016. Analysis of protein-coding genetic variation in 60,706 humans. Nature Reviews Disease Primers, 536(7616), pp.285–291.

Li, H. & Durbin, R., 2010. Fast and accurate long-read alignment with Burrows-Wheeler transform. Bioinformatics, 26(5), pp.589–595.

Liem, N.L.M. et al., 2004. Characterization of childhood acute lymphoblastic leukemia xenograft models for the preclinical evaluation of new therapies. Blood, 103(10), pp.3905–3914.

Lock, R.B. et al., 2002. The nonobese diabetic/severe combined immunodeficient (NOD/SCID) mouse model of childhood acute lymphoblastic leukemia reveals intrinsic differences in biologic characteristics at diagnosis and relapse. Blood, 99(11), pp.4100–4108.

Mayakonda, A. et al., 2018. Maftools: efficient and comprehensive analysis of somatic variants in cancer. Genome Research, 28(11), pp.1747–1756.

Peters, T.L. et al., 2015. BCOR-CCNB3 fusions are frequent in undifferentiated sarcomas of male children. Modern Pathology, 28(4), pp.575–586.

Reid, J.G. et al., 2014. Launching genomics into the cloud: deployment of Mercury, a next generation sequence analysis pipeline. BMC Bioinformatics, 15(1), p.30.

Shen, Y. et al., 2010. A SNP discovery method to assess variant allele probability from next-generation resequencing data. Genome Research, 20(2), pp.273–280.

Wang, L. et al., 2015. Genomic profiling of Sézary syndrome identifies alterations of key T cell signaling and differentiation genes. Nature Genetics, 47(12), pp.1426–1434.

Waskom, M., seaborn: statistical data visualization. Python 2.7 and 3.5. Available at: https://seaborn.pydata.org/.

Yu, L. et al., 2010. A clinically relevant orthotopic xenograft model of ependymoma that maintains the genomic signature of the primary tumor and preserves cancer stem cells in vivo. Neuro-oncology, 12(6), pp.580–594.

